# Voltage-gated calcium channels generate blastema Ca^2+^ fluxes restraining zebrafish fin regenerative outgrowth

**DOI:** 10.1101/2024.08.21.608903

**Authors:** Heather K. Le Bleu, Rea G. Kioussi, Astra L. Henner, Victor M. Lewis, Scott Stewart, Kryn Stankunas

## Abstract

Adult zebrafish fins regenerate to their original size regardless of damage extent, providing a tractable model of organ size and scale control. Gain-of-function of voltage-gated K^+^ channels expressed in fibroblast-lineage blastema cells promotes excessive fin outgrowth, leading to a long-finned phenotype. Similarly, inhibition of the Ca^2+^-dependent phosphatase calcineurin during regeneration causes dramatic fin overgrowth. However, Ca^2+^ fluxes and their potential origins from dynamic membrane voltages have not been explored or linked to fin size restoration. We used fibroblast-lineage GCaMP imaging of regenerating adult fins to identify widespread, heterogeneous Ca^2+^ transients in distal blastema cells. Membrane depolarization of isolated regenerating fin fibroblasts triggered Ca^2+^ spikes dependent on voltage-gated Ca^2+^ channel activity. Single cell transcriptomics identified the voltage-gated Ca^2+^ channels *cacna1c* (L-type channel), *cacna1ba* (N-type), and *cacna1g* (T-type) as candidate mediators of fibroblast-lineage Ca^2+^ signaling. Small molecule inhibition revealed L- and/or N-type voltage-gated Ca^2+^ channels act during regenerative outgrowth to restore fins to their original scale. Strikingly, *cacna1g* homozygous mutant zebrafish regenerated extraordinarily long fins due to prolonged outgrowth. The regenerated fins far exceeded their original length but with otherwise normal ray skeletons. Therefore, *cacna1g* mutants uniquely provide a genetic loss-of-function long-finned model that decouples developmental and regenerative fin outgrowth. Live GCaMP imaging of regenerating fins showed T-type Cacna1g channels enable Ca^2+^ dynamics in distal fibroblast-lineage blastemal mesenchyme during the outgrowth phase. We conclude “bioelectricity” for fin size control likely entirely reflects voltage-modulated Ca^2+^ dynamics in fibroblast-lineage blastemal cells that specifically and steadily decelerates outgrowth at a rate tuned to restore the original fin size.

## INTRODUCTION

Organs grow to an optimal size in scale with the individual organism. Disruption of scaling mechanisms underlies congenital disease and contributes to dysregulated, disease-associated growth. Robust organ regeneration exemplifies and extends this organ scaling mystery by the restoration of lost or damaged tissue to the original proportions. Adult zebrafish, like most teleosts, regenerate fins to the original size and pattern, providing a compelling vertebrate model to explore mechanisms of scaled organ growth.

Zebrafish fins are characterized by a series of tapered and segmented skeletal structures termed lepidotrichia or fin rays. Each fin ray segment comprises two opposing hemi-ray bones produced by ray-lining osteoblasts. The hemi-rays enclose sensory nerves, blood vessels, and fibroblast cells. Fin injury triggers wound epidermis formation and cell dedifferentiation (reviewed in Sehring and Weidinger, 2022). Lineage-restricted progenitor cells then migrate, proliferate, and organize to establish ray-associated regenerative blastemas (Knopf et al., 2011; Singh et al., 2012; Sousa et al., 2011; Stewart and Stankunas, 2012; Tu and Johnson, 2011). Distal fibroblast-derived blastemal mesenchyme transitions to a growth factor-producing “organizer” or “niche” state to initiate outgrowth around 3 days post-amputation (Wehner et al., 2014; Stewart et al., 2019). Outgrowth progressively slows until the original fin size is restored (Morgan, 1900; Iovine and Johnson, 2000; Stewart et al., 2021).

Zebrafish mutants with disrupted fin scaling provide an entry to investigate fin outgrowth control (van Eeden et al., 1996). Strikingly, all such mutants implicate ion signaling. *shortfin* (*sof*) fish with disrupted Gap junction alpha-1 protein (Connexin-43) develop small fins with shortened bony ray segments (Iovine et al., 2005). Long-finned zebrafish genetic models nearly all reflect gain-of-function of potassium K^+^ channels. *another longfin* (*alf^ty86d^*) fish develop and regenerate long fins due to a gain-of-function mutation in the K^+^ channel *kcnk5b* (Daane et al., 2018; Perathoner et al, 2014). *longfin^t2^* (*lof^t2^*) fin overgrowth is caused by ectopic expression of *kcnh2a,* an ether-a-go-go (EAG)-related voltage-gated K^+^ channel in fin fibroblast-lineage cells (Stewart et al., 2021; Daane et al., 2021). Further, transgenic overexpression of K^+^ channels *kcnj13*, *kcnj1b*, *kcnj10a*, and *kcnk9* all lead to overgrown fins (Silic et al., 2020). Roles for gap junctions and voltage-gated, membrane potential-setting K^+^ channels suggest “bioelectricity” patterns and/or modulates the extent of fin outgrowth. However, mechanisms linking ion channels and altered membrane potential dynamics to outgrowth-determining processes are unclear.

Calcineurin inhibition during fin regeneration leads to dramatically overgrown fins similar to genetic long-finned models (Kujawski et al., 2014; Stewart et al., 2021). Calcineurin, a Ca^2+^ dependent protein phosphatase, likely is an ion signaling node for fin outgrowth control, acting upstream (Daane et al., 2018; Yi et al., 2021) and/or downstream (Stewart et al., 2021) of K^+^ channel-modulated membrane potentials and, presumably, Ca^2+^ dynamics. Calcineurin activity exclusively acts during the outgrowth phase to gradually slow and eventually halt growth as fin size is restored (Stewart et al., 2021). Likewise, fibroblast-lineage ectopic Kcnh2a in *longfin^t2^*mutants acts uniquely during outgrowth to drive regenerating fin overgrowth (Stewart et al., 2021). However, Ca^2+^ fluxes and their control, including by Ca^2+^ channels, in fibroblast-lineage cells of regenerating fins have not been examined.

We explored if membrane potential-regulated Ca^2+^ signaling in fibroblast-lineage blastemal cells of regenerating fins contributes to growth cessation and organ size restoration. We generated a GCaMP6s reporter line expressed in regenerating fin fibroblasts to determine that blastemal cells flux Ca^2+^ in vivo and respond to membrane depolarization dependent on voltage-gated Ca^2+^ channels in vitro. Single cell transcriptomics and *in situ* expression studies localized specific L-, N-, and T-type voltage-gated Ca^2+^ channels to fibroblast-lineage blastema mesenchyme during fin regenerative outgrowth. Pharmacological inhibition showed L- and/or N-type voltage-gated Ca^2+^ channels restrain regenerating fin outgrowth in vivo. T-type *cacna1g* channel mutants exhibited strikingly out-of-scale regenerative overgrowth, providing a unique loss-of-function long-finned model. Live imaging of regenerating adult fins demonstrated that Cacna1g enables Ca^2+^ fluxes in fibroblast-lineage blastema cells, specifically tying the Ca^2+^ dynamics to outgrowth control. We conclude voltage-gated Ca^2+^ channel activity within fibroblast-derived blastemal mesenchyme produces a tuned rate of decelerating outgrowth that restores scaled fin size.

## RESULTS

### Voltage-gated Ca^2+^ channels mediate Ca^2+^ dynamics within fibroblast-lineage regenerating fin cells

K^+^ channel gain-of-function and calcineurin inhibition likely disrupt the same ion signaling mechanism to produce long-finned zebrafish (Perathoner et al., 2014; Stewart et al., 2021), suggesting Ca^2+^ as a second messenger moderating fin outgrowth. Further, ion signaling likely acts within fibroblast-lineage cells to gradually terminate fin regenerative outgrowth (Daane et al., 2021; Stewart et al., 2021). We generated a transgenic line using the *tryptophan hydroxylase 1b* (*tph1b*) promoter (Kapsimali et al., 2011) driving GCaMP6s (Chen et al., 2013) expression to monitor Ca^2+^ dynamics within fibroblast-lineage cells of regenerating fins (Stewart et al., 2021; Tornini et al., 2016). 3 day post-amputation (dpa) *tph1b:GCaMP6s* regenerating fins showed GCaMP6 fluorescence specifically in blastemal tissue extending from each ray (Figure 1A). Whole mount antibody staining confirmed GCaMP6s expression in fibroblast-lineage distal blastema cells by co-expression of the Dachshund (Dach) transcription factor (Lewis et al., 2023; Stewart et al., 2021) (Figure 1B). We imaged GCaMP6s dynamics in 3 dpa regenerating fins of live zebrafish by spinning disc confocal microscopy and then measured individual cell volumetric GCaMP6s fluorescence over time. Many blastemal cells exhibited fluctuating GCaMP6s levels, including spikes of variable duration and oscillatory pattern (Supplemental Movie 1; Figure 1C, D). We conclude fibroblast-lineage distal blastema cells flux Ca^2+^ levels in vivo and hypothesize such dynamics represent ion signaling modulation of the fin outgrowth period and therefore fin size.

**Figure 1.**
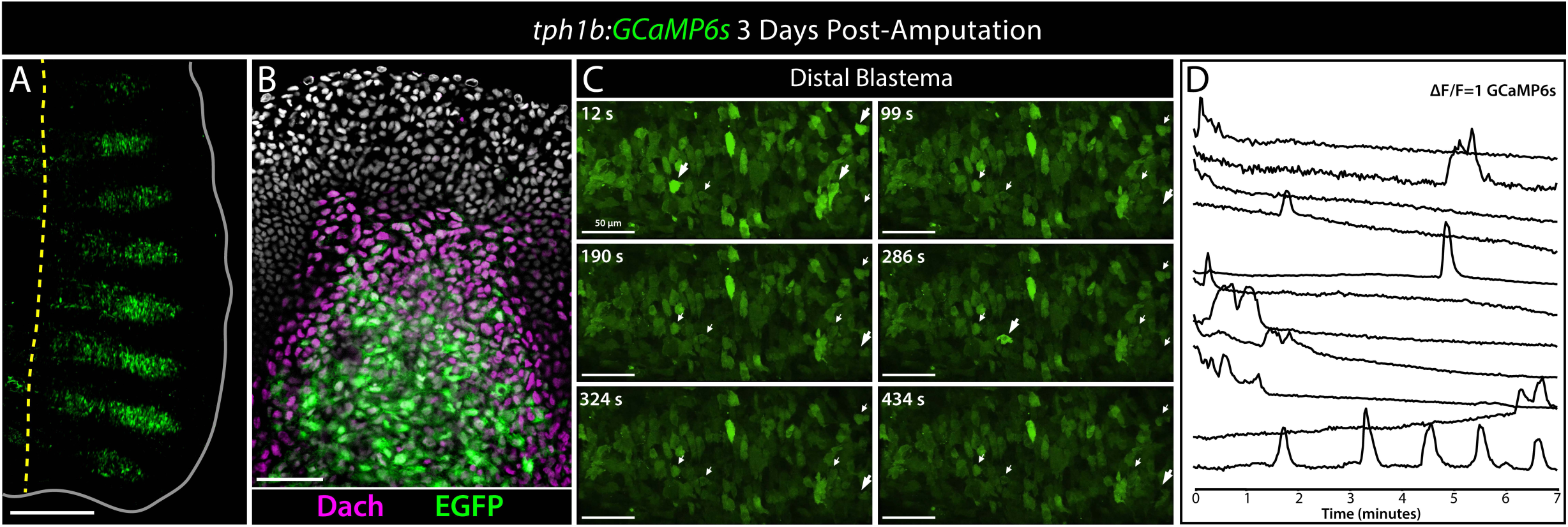
The *tph1b:GCaMP6s* line demonstrates Ca^2+^ fluxes in distal blastemal cells of regenerating fins. **(A)** Whole mount confocal image of a 3 day post amputation (dpa) regenerating caudal fin of a *tph1b:GCaMP6s* (green) adult fish. The yellow dashed line shows the amputation plane. The fin is outlined. The scale bar is 500 µm. **(B)** *tph1b*-driven GCaMP6s expression includes distal blastemal cells. Confocal maximum intensity projection image of an antibody-stained 3 dpa *tph1b:GCaMP6s* regenerating fin section. Dachshund (Dach) and EGFP (for GCaMP6s) are in magenta and green, respectively. Hoechst-stained nuclei are gray. **(C)** Representative frames from a confocal movie showing the distal blastema of a *tph1b:GCaMP6s* regenerating caudal fin at 3 dpa. Scale bars are 50 µm. The arrows mark select cells with dynamic GCaMP6s fluorescence over the 7-minute period. The arrows are enlarged during GCaMP6s-reported Ca^2+^ spikes. **(D)** Plots of normalized GCaMP6s intensity over time for the marked cells in **(C)** showing heterogenous Ca^2+^ dynamics over the 7-minute period.

We next aimed to determine if membrane potential dynamics and voltage-gated Ca^2+^ channels contribute to blastema Ca^2+^ dynamics. Time-lapse imaging of *tph1b:GCaMP6s* primary cells prepared from regenerating fins revealed sporadic Ca^2+^ fluxes (Supplemental Movie 2). Voltage-gated Ca^2+^ channels are activated by membrane depolarization and facilitate the flow of extracellular Ca^2+^ into the cytoplasm (Catterall, 2011). We depolarized *tph1b:GCaMP6s* primary cells with KCl to investigate if membrane voltage dynamics can trigger Ca^2+^ fluxes. 56% of *tph1b:GCaMP6s*-responsive cells spiked GCaMP6s fluorescence upon membrane depolarization (Figure 2A, B; Supplemental Movie 2). The Ca^2+^ channel blockers amlodipine (L-type) (Nayler and Gu, 1991), cilnidipine (L/N-type) (Yamaura et al., 1986), and PD1732122 (N-type) (Hu et al., 1999) all reduced the fraction of excitable cells (amlodipine: 12%, 15/127 cells; cilnidipine: 4%, 4/103 cells; PD17132: 20%, 29/147 cells) compared to DMSO-treated control cells (39%, 70/109 cells; Figure 2C-F). Therefore, fibroblast-lineage regenerating fin cells have excitable characteristics with at least L- and N-type voltage-gated Ca^2+^ channels contributing to membrane voltage-regulated Ca^2+^ transients.

**Figure 2.**
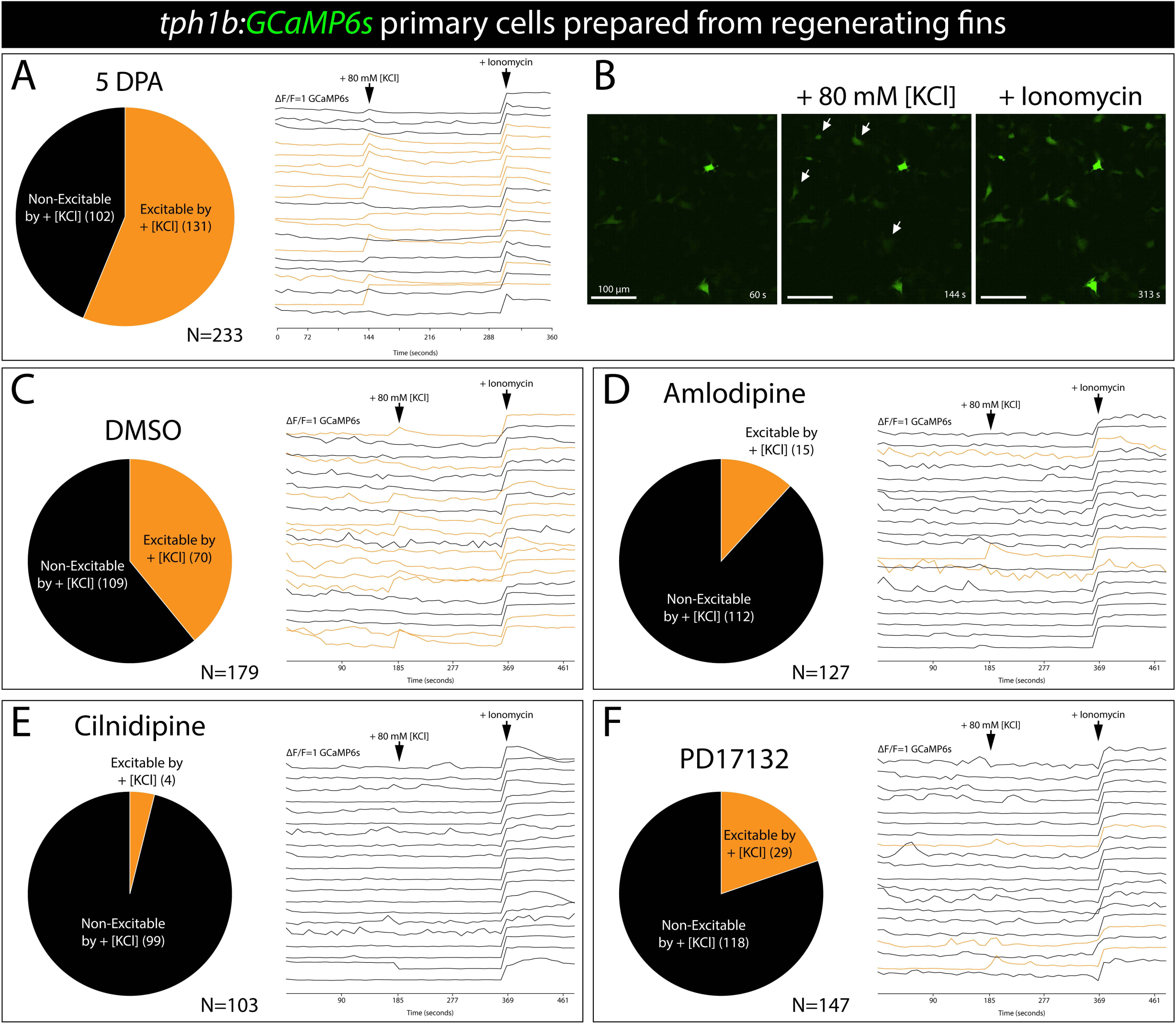
Voltage-gated Ca^2+^ channels produce Ca^2+^ fluxes upon membrane depolarization of fibroblast-lineage fin cells. **(A, B)** Fibroblast-lineage blastemal mesenchymal cells flux Ca^2+^ in response to depolarization. Primary cells were prepared from adult *tph1b:GCaMP6s* reporter fish caudal fins at 5 days post amputation (dpa). **(A)** The pie chart indicates the fraction of *tph1b:GCaMP6s*-expressing primary cells excited by addition of 80 mM KCl at 144 seconds(s). Treatment with the calcium ionophore ionomycin at 313 s demonstrates each cell’s maximum GCaMP6s signal. The neighboring plot shows normalized GCaMP6s intensity traces for twenty randomly selected KCl-responsive cells. **(B)** Representative time-lapse confocal images showing Ca^2+^ spikes at 144 s (+ 80 mM KCl) and 313 s (+ Ionomycin). White arrows point to select Ca^2+^-responsive cells. Scale bars are 100 µm. **(C-F)** L- and N-type voltage-gated Ca^2+^ channels enable calcium fluxes upon membrane depolarization of regenerating fin cells. Pie charts indicate the fraction of excitable cells upon DMSO (**C**; control), 500 nM amlodipine (**D**; L-type inhibitor), 500 nM cilnidipine (**E**, N- and L-type inhibitor), and 500 nM PD17132 (**F**, N-type inhibitor) treatment. The adjacent plots show normalized GCaMP6s intensity traces corresponding to twenty randomly selected cells. Individual Ca^2+^ traces indicate KCl-responsive (orange) and non-responsive (black) cells.

### Fin fibroblast-lineage cells express voltage-gated Ca^2+^ channels

The *longfin^t2^* (ectopic Kcnh2a) and inhibited calcineurin long-finned models disrupt ion signaling that continuously acts after 5 dpa to slow and eventually terminate outgrowth (Stewart et al., 2021). We re-analyzed single cell RNA-Seq data collected from 7 dpa regenerating caudal fins using an expanded transcriptome to distinguish voltage-gated Ca^2+^ channels expressed in fibroblast-lineage blastemal fin cells during fin outgrowth (Lawson et al., 2020; Lewis et al., 2023). We identified 10 clusters representing all major fin lineages, including co-clustered fibroblast- and osteoblast-lineage cells, annotated by established marker genes enriched within each cluster (Cao et al., 2019; Farnsworth et al., 2020; Hou et al., 2020; Lewis et al., 2023) (Figure 3A-D; Supplemental Figure 1). We surveyed the expression of genes encoding α_1_ subunit pores of L-, P-, N-, and T-type Ca^2+^ channels (Catterall, 2011) (Supplemental Figure 2). Of all candidates, L-type *cacna1c*, N-type *cacna1ba*, and T-type *cacna1g* transcripts were expressed specifically within the fibroblast/osteoblast cluster (Figure 3E-G). *cacna1g* and *cacna1c* expression was relatively broad across the cluster, including maturing fibroblasts (*rgs5a*- and *fhl1a*-expressing; Lewis et al. 2023) (Figure 3C, E, and G; Supplemental Figure 2). In contrast, *cacna1ba* was enriched in presumed distal blastema mesenchyme based on overlapping expression with *msx3*, *dachc*, *wnt5a*, and *robo3* (Lewis et al., 2023; Figure 3B and E; Supplemental Figure 2).

**Figure 3.**
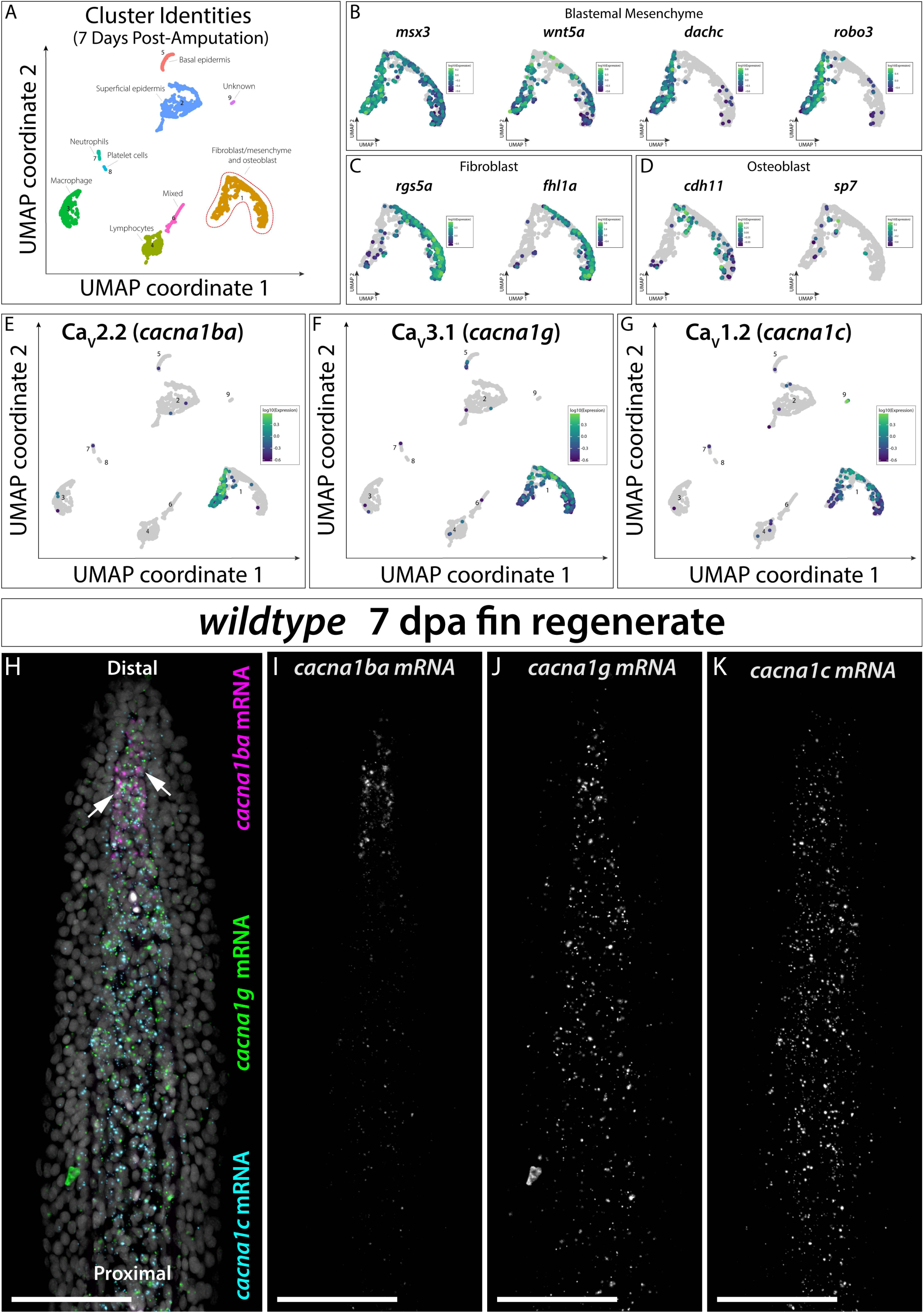
L-, N-, and T-type voltage Ca^2+^ channels are expressed in fibroblast-lineage blastemal cells. **(A)** Uniform Manifold Approximation and Projection (UMAP) showing cells from 7 day post amputation (dpa) scRNA-seq caudal fin tissue projected in 9 clusters annotated using established markers (see Supplemental Figure 1) (Lewis et al., 2023). The red dashed line outlines the cluster containing fibroblast- and osteoblast-lineage cells. (**B-D**) UMAPs of cluster 3 highlighting the indicated transcripts with known, elevated expression in **(B)** blastemal mesenchyme, **(C)** fibroblasts, and **(D)** osteoblasts. **(E-G)** UMAPs showing specific expression of the α_1_ subunit voltage-gated Ca^2+^ channels genes *cacna1ba* (**E**; N-type), *cacna1g* (**F**; T-type), and *cacna1c* (**G**; L-type) in the fibroblast/osteoblast cluster. **(H-K)** 7 dpa longitudinal section of regenerating caudal fin stained by RNAscope for *cacna1ba* (magenta), *cacna1g* (green), and *cacna1c* (cyan) transcripts. The white arrows point to *cacna1ba*, *cacna1c*, and *cacna1g* co-expressing distal blastema cells. Hoechst-stained nuclei are marked in gray. Individual max intensity projections for *cacna1ba* (I), *cacna1g* (J), and *cacna1c* (K) mRNA are shown in grayscale. Scale bars are 50 µm.

We assessed the in situ spatial distributions of *cacna1c*, *cacna1ba*, and *cacna1g* transcripts by RNAscope (Wang et al., 2012) on 7 days post-amputation (dpa) fin sections (Figure 3H-K). As predicted by the scRNA-seq analysis, *cacna1ba* exclusively was expressed in *dachc* co-expressing distal blastema (Lewis et al., 2023; Stewart et al., 2019) (Figure 3H and I; Supplemental Figure 3; Supplemental Figure 4). *cacna1c* and *cacna1g* transcripts were enriched in all fibroblast-lineage cells and adjacent blastema-lining osteoblasts (Figure 3H, J, and K; Supplemental Figure 4). We conclude these voltage-gated Ca^2+^ channels likely contribute to the observed cytosolic Ca^2+^ dynamics within intra-ray fibroblast-lineage cells.

### L/N-type Ca^2+^ channels restrain fin outgrowth during regeneration

We hypothesized voltage-gated Ca^2+^ channel activity, like calcineurin (Daane et al. 2018; Stewart et al. 2021) and ectopic Kcnh2a in *longfin^t2^* fish (Stewart et al., 2021), acts during the outgrowth phase to modulate regenerated fin size. We tested this by treating fish with cilnidipine, the dual N/L-type Ca^2+^ channel blocker most potent in the in vitro assays, during fin regeneration. Daily 5 µM cilnidipine from 5-28 dpa caused significant fin overgrowth without other apparent physiologic effects (Figure 4A-E; Supplemental Figure 5). Regenerated fins of cilnidipine-treated fish also had fewer ray segments, as with calcineurin inhibition and *another longfin* (*alf*) long-finned models (Kujawski et al., 2014; Perathoner et al., 2014) (Figure 4F-H; Supplemental Figure 5). We conclude L-and/or N-type Ca^2+^ channels actively promote fin outgrowth cessation, consistent with *cacna1c* and *cacna1ba* expression in Ca^2+^-fluxing and outgrowth-regulating blastema mesenchyme.

**Figure 4.**
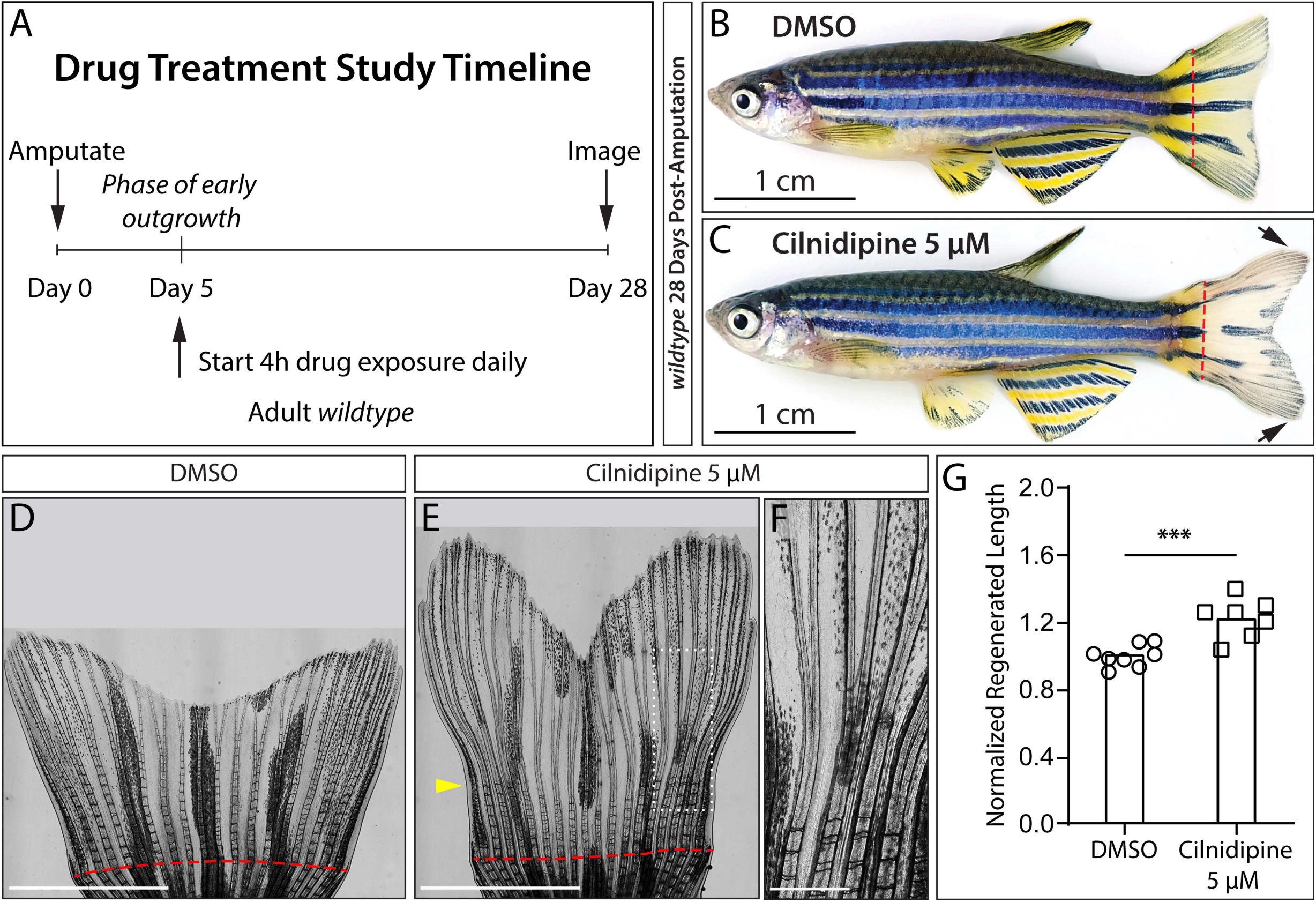
L-and/or N-type Ca^2+^ channels actively promote fin outgrowth cessation. **(A)** Timeline for outgrowth phase drug treatments of caudal fin-resected adult zebrafish from 5-28 days post amputation (dpa). **(B, C)** Representative brightfield images of **(B)** DMSO and **(C)** 5 µM cilnidipine-treated zebrafish at 28 dpa. The black arrows indicate overgrown regenerating caudal fin tissue. Dashed red lines mark amputation planes. Scale bars are 1 cm. **(D-F)** Stitched brightfield images showing 28 dpa regenerated caudal fins of the same fish in **(B)** and **(C)**. Scale bars are 4 mm. A red dashed line indicates the amputation plane. The yellow arrowhead indicates the onset of joint defects upon cilnidipine treatment initiation at 5 dpa. **(F)** The zoomed region marked by a dashed white box in **(E)**. The scale bar is 200 µm. **(G)** Graph showing body length-normalized regenerated fin lengths (measured at the 3^rd^ ray) of DMSO (control, n=8) and 5 μM cilnidipine (n=7) treated fish. Each point represents an individual fish. *\*\*\**: *P < 0.001* by an unpaired Student’s t-test.

### The T-type *cacna1g* channel centrally restricts fin outgrowth

The prominent *cacna1g* expression fibroblast-lineage blastema mesenchyme also implicates low-voltage-regulated T-type channel activity in outgrowth control. Therefore, we used CRISPR/Cas9 technology to generate loss-of-function alleles of the α_1_ subunit gene *cacna1g*. We induced deletions in the 5’-UTR and/or exon 3 coding sequence. F0 *cacna1g* crispants displayed mosaic developmental and, more dramatically, regenerative overgrowth across all fins (Supplemental Figures 6 and 7). We outcrossed founders to identify two putative *cacna1g* loss-of-function alleles. The *cacna1g^b1477^* allele had a 6 bp deletion in the 5’-UTR and a 4 bp deletion in exon 3 leading to a frame shift and early stop codon (Supplemental Figure 6). The second allele, *cacna1g^b1478^*, had a 4 bp deletion and 36 bp insertion at the exon 3 guide site, creating a frame shift into the same early stop codon (Supplemental Figure 8). Homozygous *cacna1g^b1477/b1477^*zebrafish developed modestly overgrown fins with slightly longer ray segments but appeared otherwise normal (Supplemental Figure 6). Strikingly, both *cacna1g^b1477/b1477^* and *cacna1g^b1478/b1478^* fish regenerated exceptionally long caudal fins that greatly exceeded their developed length by 24 dpa (Figure 5A-G; Supplemental Figures 6, 8, and 9). *cacna1g^b1477/+^* heterozygotes displayed only modest regenerative caudal fin overgrowth (Supplemental Figure 10). Therefore, *cacna1g^b1477^*and *cacna1g^b1478^* are most likely recessive loss-of-function alleles, thereby uncovering the requirement for voltage-gated ion signaling and specifically T-type Ca^2+^ channels in regenerated fin size restoration.

**Figure 5.**
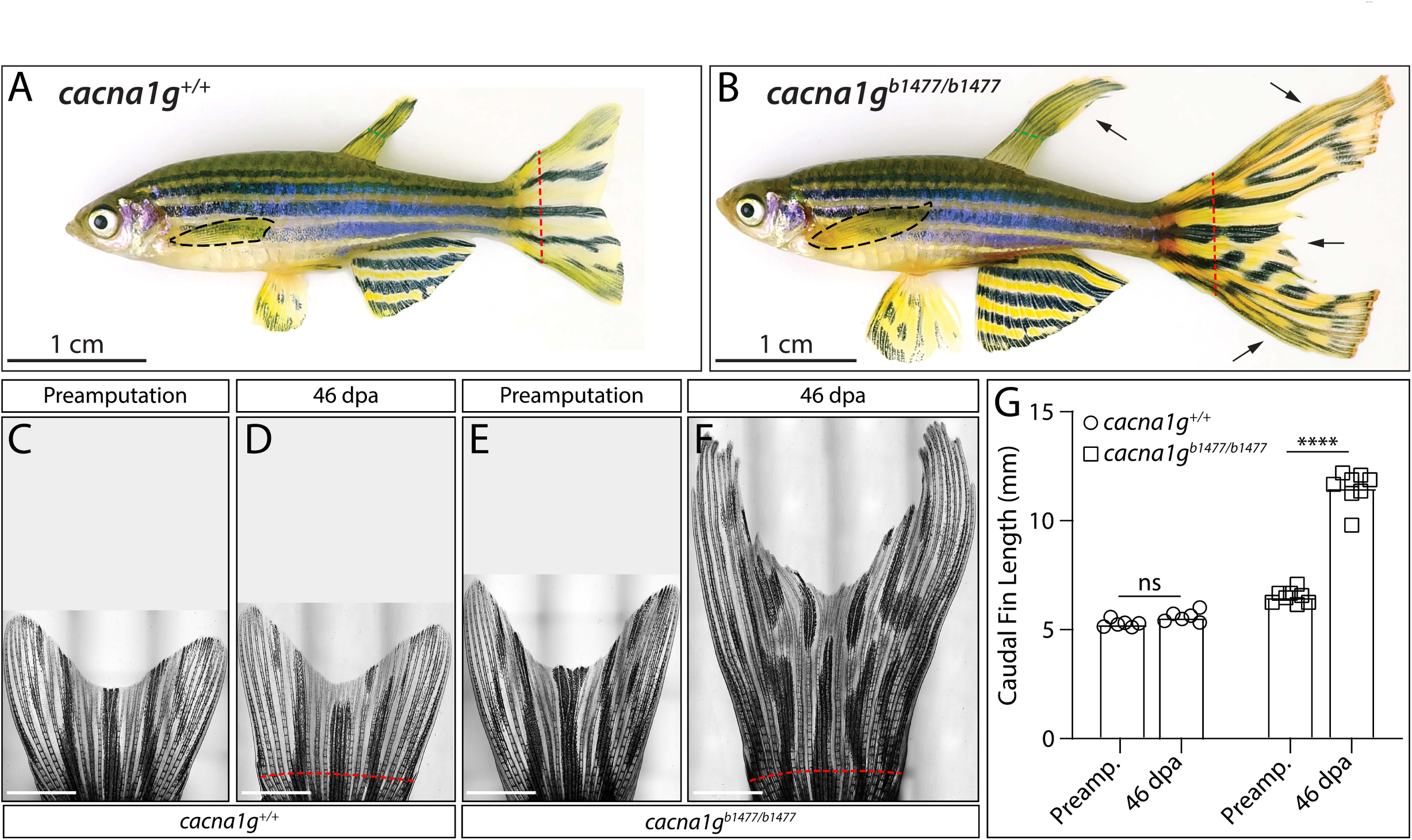
Long-finned *cacna1g* loss-of-function mutants link T-type voltage-gated Ca^2+^ channels to regenerated fin size restoration. **(A, B)** Brightfield images of representative (A) *cacna1g^+/+^* and (B) *cacna1g^b1477/b1477^* adult zebrafish including their regenerated caudal fins at 66 days post amputation (dpa). Black dashed lines outline pectoral fins. Green dashed lines indicate where dorsal fins were clipped for genotyping. Red dashed lines indicate caudal fin amputation sites. Scale bars are 1 cm. **(C-F)** Brightfield stitched caudal fin images of *cacna1g^+/+^* and *cacna1g^b1477/b1477^*fish before amputation and at 46 dpa. Red dashed lines mark amputation positions. Scale bars are 2 mm. **(G)** Graph showing caudal fin lengths (third fin ray, from the amputation plane to the fin tip) of *cacna1g^+/+^*(circles) and *cacna1g^b1477/b1477^* (squares) fish prior to amputation (preamp.) and after regeneration (46 dpa). *\*\*\*\**: *P < 0.0001* using paired two-way ANOVA and Tukey’s post-hoc tests; ns: not significant.

We focused on the *cacna1g^b1477^* allele for further characterization. *cacna1g^b1477/b1477^* fish greatly overgrew all amputated median as well as paired fins, including only the singly resected of paired pectoral fins (Supplemental Figures 11 and 12). The degree of caudal fin regenerative overgrowth even exceeded that of *longfin^t2/+^* fish amputated in parallel (Supplemental Figure 13). Further, unlike *longfin^t2^, cacna1g^b1477/b1477^* caudal fins regenerated to a length far exceeding their original scale (Supplemental Figure 13). Joint length was only modestly increased in *cacna1g^b1477/b1477^* regenerated caudal fins, more closely resembling the *longfin^t2^* than calcineurin-inhibition or *alf* models (Supplemental Figure 14). We also observed pronounced blood vessels and blood pooling in the distal tissue of 88 dpa regenerating caudal fins of *cacna1g^b1477/b1477^* fish (Supplemental Figure 15), phenotypes associated with other long-finned models (Kujawski et al., 2014; Lanni et al., 2019). Together, *cacna1g^b1477/b1477^* uniquely establishes a recessive, loss-of-function long-finned model that centrally and specifically implicates T-type voltage-gated Ca^2+^ channels in restraining regenerative fin outgrowth.

We hypothesized Cacna1g-containing channels (Ca_V_3.1) control fin size by progressively slowing and then terminating fin outgrowth, as seen with *longfin^t2^* and calcineurin-inhibited long finned models (Stewart et al., 2021). We defined the outgrowth characteristics of *cacna1g^b1477/b1477^*and wildtype clutchmates by measuring caudal fin length over a regenerative time course (Figure 6, Supplemental Figure 10). The size of *cacna1g^+/+^* and *cacna1g^b1477/b1477^* regenerating caudal fins were similar through approximately 5 days of regeneration. Thereafter, the *cacna1g^b1477/b1477^*caudal fin outgrowth rate increasingly exceeded that of control animals. Therefore, *cacna1g^b1477/b1477^* fish regenerated progressively longer caudal fins, matching the outgrowth kinetics of *lof^t2/+^* regenerating fins (Iovine and Johnson, 2000; Stewart et al., 2021) (Figure 6A, C-H). The maximum rate of outgrowth peaked at ∼3-4 dpa for both *cacna1g^+/+^* and *cacna1g^b1477/b1477^* fish and then gradually declined, with outgrowth rate curves closely fitting one-phase exponential decay curves but with differential decay rates (Figure 6B). We conclude Cacna1g maintains a steadily decreasing outgrowth rate (deceleration rate) that helps restore fins to their original size.

**Figure 6.**
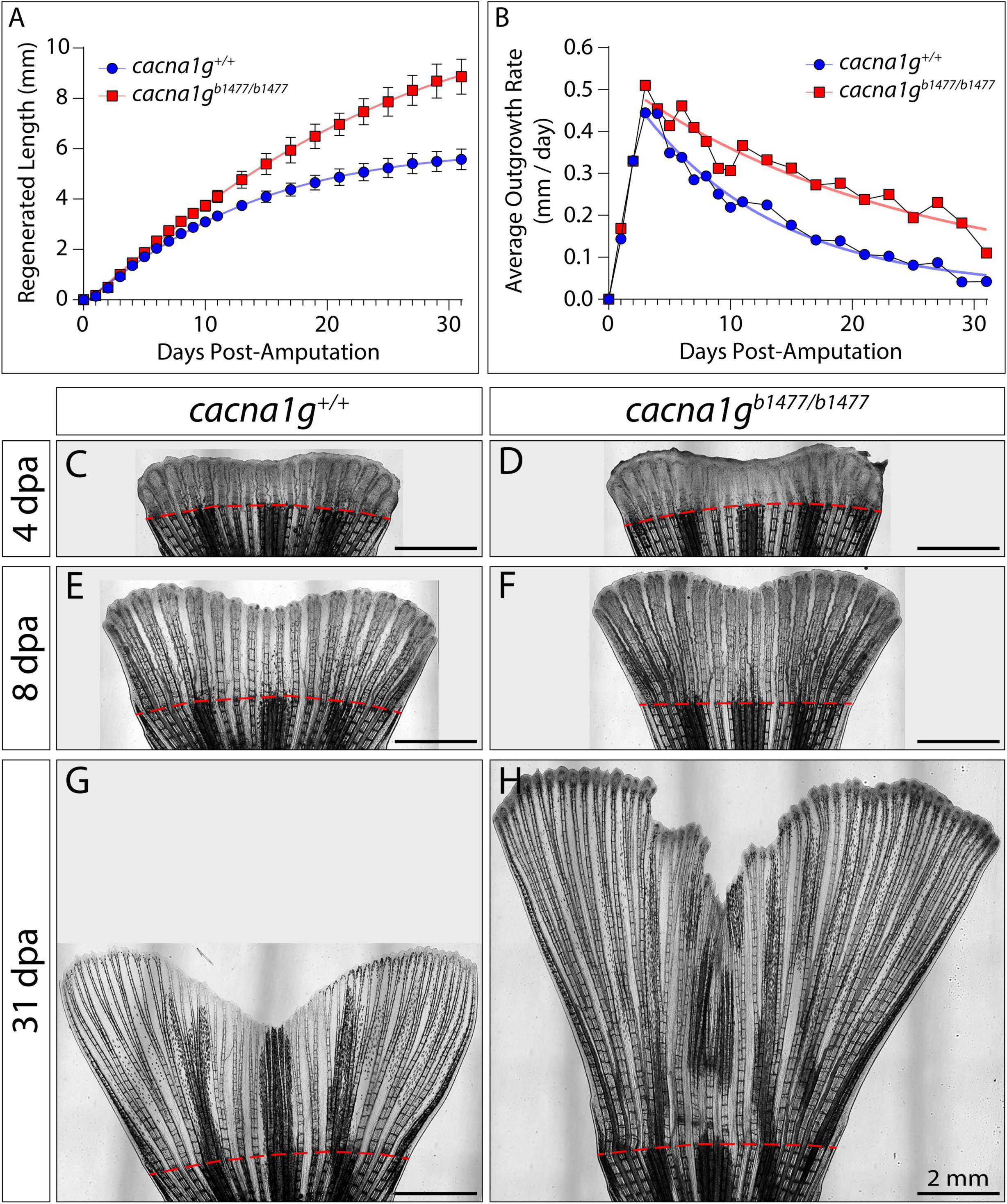
Cacna1g decelerates regenerating fin outgrowth to restore fin size. **(A, B)** Graphs showing **(A)** regenerated caudal fin lengths and derived **(B)** outgrowth rates of *cacna1g^+/+^* (blue circles) and *cacna1g^b1477/b1477^*(red squares) fish over 31 days of regeneration. **(A)** Sigmoidal logistic growth curves are fit to regenerated fin lengths over time to reflect an initial delay for blastema establishment and then a progressively slowing outgrowth rate. Means ± SD are plotted for each timepoint (n=14 for each genotype). **(B)** One phase exponential decay curves are fit to the average outgrowth rates starting at the peak outgrowth rate at 3 dpa. **(C-H)** Brightfield stitched images of 4, 8, and 31 dpa regenerated caudal fins of representative *cacna1g^+/+^*and *cacna1g^b1477/b1477^* fish. Scale bars are 2 mm.

### Cacna1g controls dynamic Ca^2+^ signaling in distal niche cells

We performed live GCaMP6s imaging of *cacna1g^b1477/b1477^*regenerating caudal fins to determine if Cacna1g influences the Ca^2+^ dynamics we observed in fibroblast-lineage blastemal mesenchyme. We observed heterogenous GCaMP6s reporter activity across distal fibroblast-lineage cells of 5 dpa wildtype regenerating fins, as seen at 3 dpa (Figure 7A, D; Supplemental Movie 3). In contrast, very few *cacna1g^b1477/b1477^* fin regenerate cells showed Ca^2+^ transients and even those cells showed only rare and modest Ca^2+^ spikes over a 5-minute time course (Figure 7B-D, Supplemental Movie 3). We conclude the Cacna1g T-type channel centrally enables Ca^2+^ dynamics in the distal fibroblast-lineage blastemal mesenchyme. These fibroblast-lineage Ca^2+^ fluxes decelerate fin outgrowth at a rate tuned to restore regenerated fin size.

**Figure 7.**
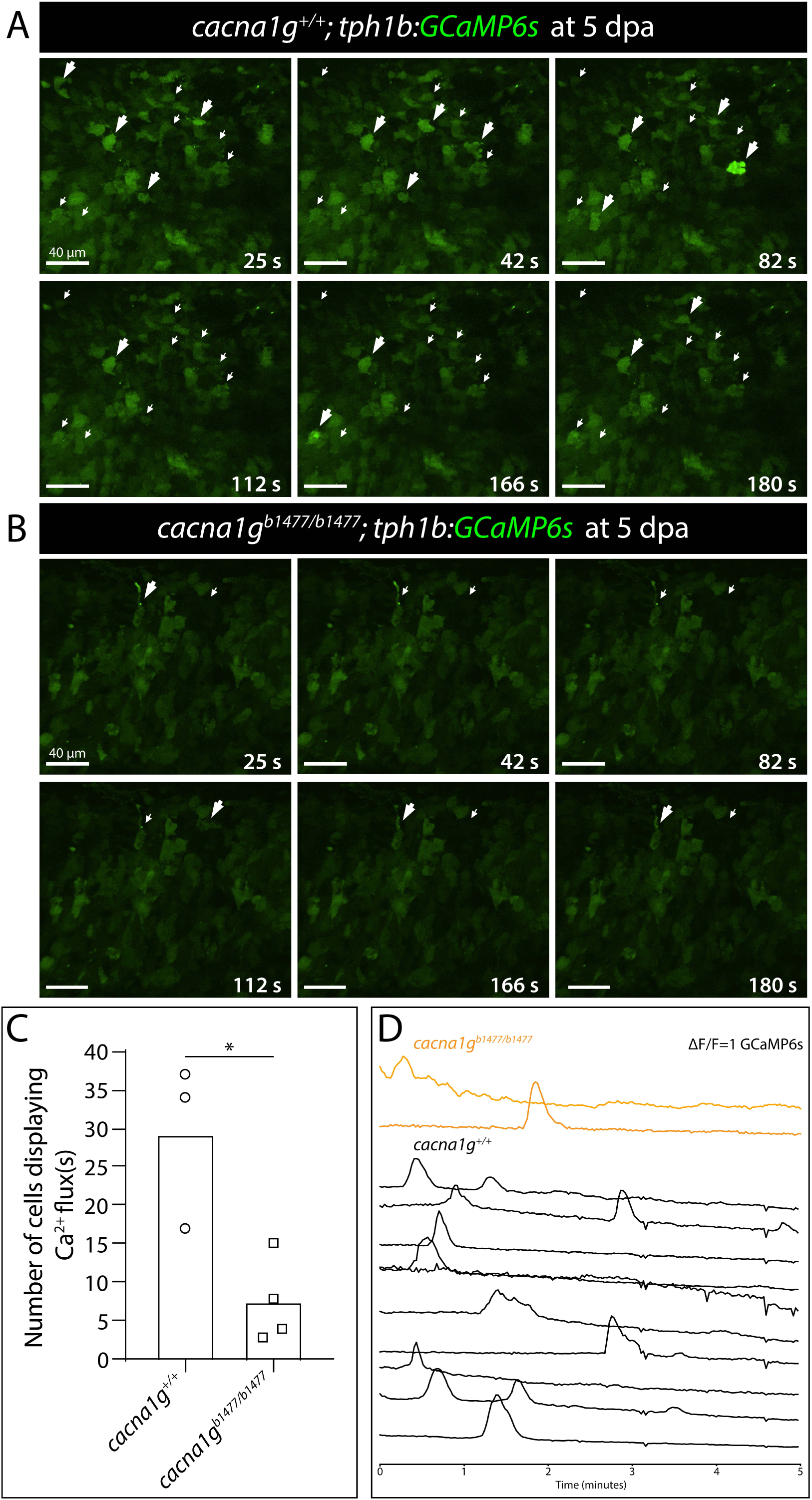
Cacnalg enables Ca^2+^ dynamics in fibroblast-derived distal blastema cells of regenerating caudal fins. **(A, B)** Confocal images from timelapse-imaged regenerating caudal fins of **(A)** *cacna1g^+/+^* and **(B)** *cacna1g^b1477/b1477^*fish expressing the *tph1b:GCaMP6s* reporter. The distal blastema is shown at 5 days post amputation (dpa). White arrows indicate Ca^2+^-fluxing cells. Arrows are enlarged when Ca^2+^ levels are elevated. Scale bars are 40 µm. **(C)** Graph showing the number of Ca^2+^-fluxing distal fibroblast-lineage blastemal cells in *cacna1g^+/+^* and *cacna1g^b1477/b1477^* 5 dpa regenerating caudal fins. Each data point represents an individual animal. *\**: *P < 0.05* by unpaired Student’s t-test. **(D)** Plots of normalized GCaMP6s intensity traces over a 5-minute period for each individual cell marked in **(A, B)**. Orange traces are *cacna1g^b1477/b1477^* cells (n=2) and black traces are *cacna1g^+/+^*cells (n=10).

## DISCUSSION

We link voltage-gated Ca^2+^ channels and downstream cytosolic Ca^2+^ fluxes within blastemal fibroblasts to the cessation of fin outgrowth. Live GCaMP6s reporter imaging of regenerating fins indicated distal fibroblast-lineage blastemal cells undergo dynamic and variable Ca^2+^ transients. Isolated regenerating fin fibroblasts spontaneously and autonomously produced heterogeneous Ca^2+^ fluxes. Further, membrane depolarization caused cytosolic Ca^2+^ spikes, indicating at least some fibroblast-lineage blastemal cells are in an “excitable” state. Inhibitor studies showed the excitability is mediated by voltage-gated Ca^2+^ channels. In vivo, fibroblast-lineage distal blastemal cells expressed *cacna1c* (L-type), *cacna1ba* (N-type), and *cacna1g* (T-type) voltage-gated Ca^2+^ channels. Temporal inhibition showed L- and/or N-type Ca^2+^ channels actively restrain fin outgrowth during late stages of regeneration. Dramatic regenerative fin overgrowth and GCaMP6s studies in *cacna1g* loss-of-function mutants revealed a central and specific role for T-type channel-dependent Ca^2+^ fluxes in fibroblast-lineage distal blastemal mesenchyme. We propose voltage-gated ion channel activity in distal fibroblast-lineage cells culminates in intracellular signaling by the classic Ca^2+^ second messenger. Ca^2+^-regulated proteins, likely the calcineurin phosphatase, then progressively restrain outgrowth and eventually terminate fin regeneration.

### Fibroblast-derived blastema cells exhibit heterogenous voltage-gated Ca^2+^ dynamics

Live-imaged regenerating caudal fins of adult *tph1b:GCaMP6s* fish showed frequent GCaMP6s activity bursts at 3 and 5 dpa, indicative of cytosolic Ca^2+^ transients. The calcium-fluxing cells were distally concentrated and therefore likely fibroblast-derived, growth factor-producing “niche”-state, or “organizing center” cells (Stewart et al., 2019; Wehner et al., 2014). The amplitude, duration and frequency of fluxes were qualitatively heterogeneous and not overtly coordinated, arguing against the transmission of bioelectric signals across a blastema field. However, coordinated calcium signaling may initiate at later outgrowth periods than we could assay (maximum 5 dpa due to loss of *tph1b*-driven GCaMP6s expression) or in distalmost blastema cells where the *tph1b:GCaMP6s* line expresses poorly.

Isolated cells prepared from regenerating *tph1b:GCaMP6s* fins showed voltage-gated Ca^2+^ channel-dependent Ca^2+^ transients, indicating cell autonomous initiation of membrane potential fluxes. Further, membrane depolarization by KCl addition spiked cytosolic Ca^2+^, again dependent on voltage-gated Ca^2+^ channels. This “excitability” as well as the GCaMP6s intensity and frequency were heterogenous, similar to in vivo observations. The variability in Ca^2+^ signaling across fibroblast-lineage cells may reflect heterogeneous cell states with differential resting membrane potential and/or responsiveness to voltage dynamics. Such differences could originate from the variable expression of voltage-gated ion channels, including *cacna1ba*, *cacna1c*, and *cacna1g*, as well as interacting ion channels and associated proteins.

### Voltage-gated Ca^2+^ channels actively restrain fin outgrowth

N- and/or L- as well as T-type voltage-gated Ca^2+^ channels contribute to the observed Ca^2+^ fluxes and restoration of fin size. The L- and N-type Ca^2+^ inhibitor, cilnidipine, displayed the most potent block of depolarization-induced Ca^2+^ flux in isolated regenerating fibroblasts. Further, cilnidipine administration to regenerating animals produced significant fin overgrowth. However, regenerating fin overgrowth was considerably more dramatic in T-type channel *cacna1g* homozygous mutants. The difference could reflect incomplete N- and/or L-type channel inhibition due to in vivo pharmacokinetics or a relatively more central role for the T-type channel in vivo.

Fibroblast-lineage blastemal mesenchyme Ca^2+^ fluxes were almost completely lost in *cacna1g^b1477/b1477^* regenerating fins. Therefore and given *cacna1g^b1477/b1477^* caudal fins otherwise regenerated near normally, fibroblast-lineage Ca^2+^ fluxes appear dedicated to outgrowth control. Further, the low-voltage T-type Cacna1g channel appears to enable fibroblast-derived blastemal mesenchyme to become voltage-responsive and/or to actively initiate voltage-gated calcium dynamics that slow outgrowth. For example, Cacna1g could respond to small membrane potential changes to produce further cation influx that then triggers high-voltage L- and/or N-type Ca^2+^ channels. These channels then produce higher amplitude and/or long duration downstream Ca^2+^ fluxes that stimulate cellular responses. Such potential cooperation between T- and L-/N-type channels could explain why in vitro inhibition of L- and N-type channels decreases the fraction of excitable cells monitored by GCaMP6s responsiveness, reminiscent of the near loss of in vivo Ca^2+^ transients in the T-type *cacna1g* mutants.

Single cell transcriptomics and in situ hybridizations of 7 dpa caudal fins revealed expression of *cacna1c* (L-type) and *cacna1g* (T-type) primarily in fibroblast-lineage blastema and osteoblasts. In contrast, *cacna1ba* (N-type) transcripts exclusively were found in far distal fibroblast-lineage cells. The overlapping expression of *cacna1ba*, *cacna1c*, and *cacna1g* in these distalmost cells could reflect where ion signaling functionally restrains outgrowth if the three channel types cooperate cell autonomously. Regardless, the upregulation of voltage-gated Ca^2+^ channels and other ion signaling transcripts in distal blastema cells appears a key gene expression component of fibroblast cell state transitions underpinning regeneration (Hoptak-Solga et al., 2008; Lewis et al., 2023; Nechiporuk & Keating 2002; Stewart et al., 2019; Wehner et al., 2014).

### A specific role for T-type Ca^2+^ channel Cacna1g and downstream cytosolic Ca^2+^ dynamics in restraining regenerative fin overgrowth

The extent of developmental fin overgrowth in *cacna1g^b1477/b1477^* fish was less than seen with either *longfin^t2^* or *alf^ty86d^*. However, after fin amputation, *cacna1g* mutant fins regenerated to extreme lengths even exceeding *longfin^t2^*. This decoupling between developed and regenerated fin length is a unique feature of the *cacna1g* long-finned model. A simple explanation why Cacna1g preferentially restrains overgrowth during fin regeneration could be functional redundancy. For example, a redundant and/or compensatory voltage-gated Ca^2+^ channel could be expressed in developing, but not regenerating, fins. Alternatively, variant outgrowth and/or scaling mechanisms could act in developing vs. regenerating fins, with only the latter requiring Cacna1g-modulated Ca^2+^ dynamics. For example, Cacna1g could be responsible for returning fin fibroblasts to an excitable state after injury without contributing to establishing said state developmentally.

Cacna1g is remarkably specific for outgrowth cessation as its loss-of-function otherwise did not overtly disrupt fin pattern, including bony ray branching and segmentation. *cacna1g^b1477/b1477^* ray segments were only slightly longer than wildtype counterparts. In contrast, cilnidipine-treated fins regenerated without ray junctions, reminiscent of *alf^ty86d^*and calcineurin-inhibited fish (Kujawski et al., 2014; Perathoner et al., 2014). This result further reinforces ray segmentation and outgrowth are orthogonal processes. Extending this idea, Ca^2+^ and calcineurin may have distinct functions in fibroblast lineage (outgrowth control) and osteoblast lineage (joint formation) cells. Perturbations that disrupt ion signaling in both cell lineages (*alf*, *sof*, FK506 treatment, cilnidipine) then produce both outgrowth and joint phenotypes. In contrast, long-finned models that disrupt ion signaling in the fibroblast but not osteoblast lineage (*longfin^t2^*, with fibroblast-specific ectopic Kcnh2a; presumably *cacna1g* homoyzgous mutants) lead to excessive outgrowth but normal segmentation. Considering *cacna1c* expression in osteoblast-lineage blastemal cells and that cilnidipine inhibits L-type channels, *cacna1c* likely contributes to cytosolic Ca^2+^ signaling for ray joint formation. Loss-of-function studies of *cacna1c* as well as the N-type *cacna1ba* channel will help resolve distinct, pathway, or redundant roles of each of the three blastema-expressed channels.

### Linking ion signaling to mechanisms of fin size restoration

A likely effector of voltage-gated Ca^2+^ channel-dependent Ca^2+^ dynamics in distal-fibroblast-derived blastema cells is the Ca^2+^-dependent phosphatase calcineurin given its pharmacologic inhibition during fin regeneration produces overgrowth (Kujawski, 2014). Further, voltage-gated Ca^2+^ channels activate calcineurin in other contexts (Cano et al., 2005; Chen et al., 2011; Gao et al., 2012; Graef et al., 1999). Gain-of-function K^+^ channels producing long-finned phenotypes then could act upstream of calcineurin by hyperpolarizing fibroblasts (as indicated for the *alf*/Kcnk5b K^+^ channel; Perathoner et al., 2014) and restricting voltage-gated Ca^2+^ channels from producing calcineurin-activating Ca^2+^ dynamics. Alternatively, calcineurin could act upstream of *alf*/Kcnk5b (Daane et al., 2018; Yi et al., 2021) to modulate other Ca^2+^-dependent signaling molecules. Finally, calcineurin could contribute to a feedback loop and thereby act both upstream and downstream of K^+^ channel-modulated ion signaling.

Our cilnidipine timing-of-administration experiments revealed that L- and/or N-type channels actively function to terminate outgrowth during fin regeneration. Similarly, temporal inhibition shows calcineurin is continuously required to drive overgrowth (Daane et al., 2018) and exclusively contributes to fin size restoration during the slowing outgrowth phase (Stewart et al., 2021). Likewise, ectopic Kcnh2a in *longfin^t2^* only drives fin overgrowth during outgrowth with no effect during the establishment phase. None of calcineurin inhibition, ectopic Kcnh2a in *longfin^t2^*, or the *cacna1g* loss-of-function mutant introduced here elevate the initial, peak fin outgrowth rate around 3-4 dpa (Stewart et al., 2021). Calcineurin inhibition also does not alter the blastema pre-pattern (Tornini et al., 2016). These results all point to a role for voltage-gated Ca^2+^/calcineurin signaling in progressively slowing the outgrowth phase rather than blastema establishment. Calcineurin also specifies the posterior edge of “hole punch” injured fins (Cao et al., 2021). However, this early tissue polarity role seems distinct from calcineurin’s role in restraining the fin outgrowth period.

We propose ion signaling or “bioelectricity” exclusively contributes to organ size of regenerated fins by promoting voltage-gated Ca^2+^ fluxes in fibroblast-lineage blastema mesenchyme during outgrowth. Calcium dynamics, likely as second messengers activating calcineurin, slow and then end outgrowth, possibly by gradually depleting the distal growth-promoting blastemal cells (Stewart et al., 2019). The underlying mechanism could include Ca^2+^/calcineurin repression of pro-progenitor growth factor production (Jiang et al., 2024; Yi et al., 2021). Regardless, by this model, bioelectricity does not produce a growth-determining pre-pattern while the blastema is established and therefore does not provide positional information itself. Instead, voltage-regulated Ca^2+^ signaling could be an effector of mechanisms “sensing” size restoration. More straightforwardly, Ca^2+/^calcineurin could maintain a robustly “tuned” countdown timer that precisely reads out positional information, which simply could be the size of the blastema population established at the onset of regeneration (Stewart et al. 2019; Wang et al., 2019).

## MATERIALS AND METHODS

### Zebrafish strains and maintenance

Zebrafish were housed in the University of Oregon Aquatic Animal Care Services facility at 28-29°C. The University of Oregon Institutional Animal Care and Use Committee oversaw animal use. Wildtype *AB* (University of Oregon Aquatic Animal Care Services) and *TL* (Haffter et al., 1996), lines were used. For all experiments, adult fish (>3 months old) of equal size and sex were used for analysis.

### Construct and generation of transgenic *tph1b:GCaMP6s* animal

The 5E_tph1b promoter plasmid was produced by PCR amplification of the zebrafish *tph1b* promoter (Kapsimali et al., 2011) and ligation into the 5E_MCS vector (Kwan et al., 2007). The 5E_tph1b promoter, ME_GCamp6s (Chen et al., 2017), 3E_polyA, and tol2_cmlcEGFP plasmids (Kwan et al., 2007) elements were combined using a Gateway LR reaction (Thermo) to generate the tph1b:GCaMP6s_polyA_cmlc2_EGFP tol2 compatible transgenesis vector. This construct was co-injected with capped RNA encoding the Tc transposase (Kawakami et al., 2000) into one stage AB embryos at a concentration of 25 ng/µl. Animals positive for EGFP expression in the heart at 48 dpa were selected, reared to adulthood, and screened for GCaMP6s expression in regenerative fin rays at 3 dpa. Founders were then outcrossed to AB fish and progeny were selected for EGFP^+^ hearts, reared to adulthood, screened, and scored for robust GCaMP6s expression in all 18 bony fin rays at 3 dpa. Multiple generational out-crossing isolated a stable *tph1b:GCaMP6s* line, which is maintained in heterozygous state by outcrossing to AB fish and selecting animals with EGFP heart expression.

### CRISPR/Cas9-generation of *cacna1g* mutants

Guide RNAs (gRNA) targeting the 5’-UTR and/or exon 3 of *cacna1g* were designed using ChopChopV3 (Labun et al., 2019). gRNA DNA templates were synthesized following a published protocol (Bassett et. al, 2013). gRNA quality was assessed prior to injection into one-stage *AB wildtype* embryos at a concentration of 100 ng/µl gRNA(s) and 500 ng/µl Cas9 protein (Thermo Fisher). F0 fish were screened for fin outgrowth in uninjured and injured states and then outcrossed to wildtype AB fish. F1 clutches harboring unique mutant alleles were identified by isolating and Sanger sequencing DNA and cDNA from fin clips, then reared to adulthood and outcrossed to AB. Mutations in 5’ - UTR were identified by amplifying genomic DNA using *cacna1g*_sg1_for and *cacna1g*_sg1_rev primers (Bhattacharya and Van Meir, 2019). cDNA was amplified from 4 dpa *cacna1g* mutant fins using *cacna1g*_sg2_for and *cacna1g*_sg2_rev primers and sequenced to identify mutations in exon 3.

### Adult whole zebrafish and fin imaging

Whole animal images were captured using a homemade light box made from a fenestrated styrofoam container, an AmScope LED-8WD lamp, and a Fujifilm X-A1 camera with a Fujinon 28mm 1.4R lens. High resolution fin images were obtained from tricaine-euthanized adult fish mounted with water on a glass slide. Stitched differential interference contrast (DIC) images were captured using a motorized Nikon Eclipse Ti-E widefield microscope with a 4X objective and NIS-Elements software.

### Single cell RNA-sequencing data analysis

Single cell RNA-Seq data (Lewis et al., 2023) was processed using the 10X Genomics Cell Ranger pipeline (version 5.0.1) and zebrafish reference genome GRCz11_104. To accommodate scRNA-Seq 3’ end-sequencing bias, reads were mapped to the comprehensive zebrafish transcriptome V4.3.2 (Lawson et al., 2020). Cell Ranger output files (barcodes, genes, and matrix files) were loaded into Monocle3 (1.0.0) for pre-processing, visualization, and clustering using R (version 4.1.2). The UMI count threshold was set at the 95th percentile to remove potential doublets. Cells expressing fewer than 250 genes also were excluded. 7,250 cells with a median of 483 genes per cell were used for downstream data analysis. Dimensionality reduction used PCA on log transformed expression matrix with 11 dimensions applied. Uniform Manifold Approximation and Projection (UMAP) dimensionality projection then grouped similar cells according to global expression profiles in 2 dimensions (Becht et al., 2018; Blondel et al., 2008; McInnes et al., 2018). The UMAP parameters were set to: metric = cosine, distance = 0.1, and neighbor = 100. Clustering using default parameters (except for cluster_cells: method = ‘louvain’, res = 1e^-6^), produced 10 clusters. Genes defining each cluster were identified by calling the top_markers() function in Monocle3 using default parameters (except for group_cells_by=”cluster”, reference_cells=1000, top_n(6)) (Cao et al., 2019). Cluster identities were assigned using these markers and genes with established expression in known regenerating fin cell types. Red blood cells were removed and UMAP clustering re-performed and annotated as shown in Supplemental Figure 1A.

### Small molecule treatments

Cilnidipine (Cayman Chemicals) was dissolved in DMSO (Sigma) and diluted to a working stock of 50 mM. From 5 days post-amputation onwards, fish were treated 4 hours a day with DMSO (0.01%) or the indicated concentration of cilnidipine and then returned to normal water flow conditions. For in vitro experiments, compounds were dissolved in DMSO and added to cell culture media in chamber slides at least 15 minutes prior to time-lapse imaging using a stage-mounted incubation system on a Nikon Ti-E inverted microscope with a Yokogawa CSU-W1 spinning disk confocal unit.

### Size measurements

All fin lengths and segments were measured along the third ray using stereomicroscope images and FIJI software (NIH). Fin length measurements started from the position of the first procurrent ray to the tip of the fin in both injured and uninjured animals. Standard body length was measured from the tip of the mouth to the caudal peduncle.

### RNAscope staining

RNAscope probes to detect *cacna1ba*, *cacna1c*, *cacna1g*, *and dachc* mRNA were designed and synthesized by ACD Bio. RNAscope was performed using the Multiplex Fluorescent kit (ACD Bio) according to manufacturer’s recommendations for paraffin embedded sections with minor modifications. Nuclei were visualized by Hoechst staining (Thermo Fischer). Imaging used Zeiss LSM 880 or LSM 710 laser scanning confocal microscopes.

### Calcium imaging and analysis

For *in vivo* fin regeneration studies, fish were anesthetized in 0.02% tricaine methanesulfonate (MS-222, Syndel) and mounted with fish water on the bottom of a chambered cover glass system (C1-1.5H-N, Cellvis). Fins were positioned with a single-hair paint brush and distal ventral rays imaged using a Nikon Ti-E/Yokogawa CSU-W1 spinning disk confocal microscope equipped with a 40X water immersion objective lens and using NIS-Elements software. The distance between confocal planes was set at 2.5 µm for Z-stack, and a time-lapse of 5 to 6 Z-stacks was conducted with an interval of 1 second between each stack for duration of 10 minutes. Post time-lapse imaging, fish were returned to tanks to recover to normal swimming behavior. Time-lapse recording of regenerated fins was processed for data acquisition using Imaris (v9.9.1). Videos were drift corrected and 3D surfaces manually drawn onto cells of interest. Each cell’s average GCaMP6s fluorescence across all frames was measured and outputted in a spreadsheet. Python scripts were used to normalize raw GCaMP6s trace data using the formula: ΔF/F = (F_t_ – F_min_)/ F(_max_ – F_min_), where F_min_ and F_max_ are the maximum and minimum trace values, respectfully (Ho et al., 2021).

For *in vitro* studies, regenerated caudal fins from *tph1b:GCaMP6s* fish were collected, and cells isolated as previously described (Stewart et al., 2014) with minor modifications. Cells were plated on chambered glass coverslips and cultured overnight at 30°C in an air incubator in L-15 media (Thermo Fisher) containing 10% Fetal Bovine Serum (Sigma). The following day, cells were equilibrated in Tyrode’s solution (10 mM HEPES, 127 mM NaCl, 12 mM NaHCO_3_, 5 mM KCl, 5 mM glucose, 1 mM MgCl_2;_ Boston Bioproducts). 20 minutes prior to imaging, cells were pretreated with the indicated small molecules or DMSO vehicle. Drug concentrations were maintained throughout the experiment at the following final concentrations: 500 nM cilnidipine; 500 nM FK506; 5 μM amlodipine; 5 μM ML218; 2 μM PD17132. Cells were imaged on a spinning disk confocal microscope for 5 minutes, depolarized by adding Tyrode’s solution containing KCl (80 mM final concentration) and imaged for an additional 5 minutes. Ionomycin (1 μM final concentration) was then added followed by 1 minute of further imaging. GCaMP6s fluorescence trace data was normalized to the highest signal timepoint (=1). Calcium-responding cells were defined as cells that reached a maximum GCaMP6s fluorescent intensity of >= 0.9 after ionomycin addition. From these filtered responding cells, the fraction of “excitable” cells susceptible to depolarization at a final concentration of 80 mM KCl was identified. Excitable cells were defined as those cells showing a normalized GCaMP6s fluorescent intensity >= 0.25 immediately after KCl administration. Twenty such Ca^2+^ fluxing cells were randomly sampled and plotted for each condition.

### Statistical analyses

Fin outgrowth analyses used paired or unpaired Student’s T-tests or one- or two-way ANOVA. For ANOVA, Tukey’s post-hoc tests were used to determine statistically significant differences between groups. A sigmoidal logistic curve was fit to regenerated ray length over time to account for the establishment phase of blastema formation. A one phase exponential decay curve was fit to outgrowth rate over time starting with the 3 dpa timepoint to isolate the outgrowth phase. All summary statistics, including mean, standard deviation and/or sample size are described in the figure legends. All calculations were performed using GraphPad Prism Version 8.4.3.

### Primers

The following genotyping primers were used:

*cacna1g*_sg1_for 5’ AGGCGTGTATTGGAAGTTGAAT 3’

*cacna1g*_sg1_rev 5’ AAGCAGTCCAAATGATGGTTCT 3’

*cacna1g*_sg2_for 5’ GAGATACCCATAAACCTGCTCG 3’

*cacna1g*_sg2_rev 5’ GGCTGGTAGGTAGGGAGAAGTT 3’

## ACKNOWLEDGEMENTS

We thank the University of Oregon Aquatic Animal Care Services for animal husbandry, Drs. Marika Kapsimali and Lilianna Solnica-Krezel for reagents, the University of Oregon Genomics and Cell Characterization Core Facility for imaging support, and Pete Batzel for advice on scRNA-Seq analysis.

## Author contributions

H. K. L., S. S., and K. S. designed experiments. H. K. L., R. G. K., A. H. L., S. S., and K. S. performed experiments. V. M. L. assisted with bioinformatics. H. K. L., S. S. and K. S. prepared and wrote the manuscript with input from all the authors.

## Data and material availability

Requests for materials should be addressed to K. S.

