## Supplementary Information for "Voltage-gated calcium channels generate blastema Ca^2+^ fluxes restraining zebrafish fin regenerative outgrowth"

**Figures S1-S15**

**Movies S1-S3**

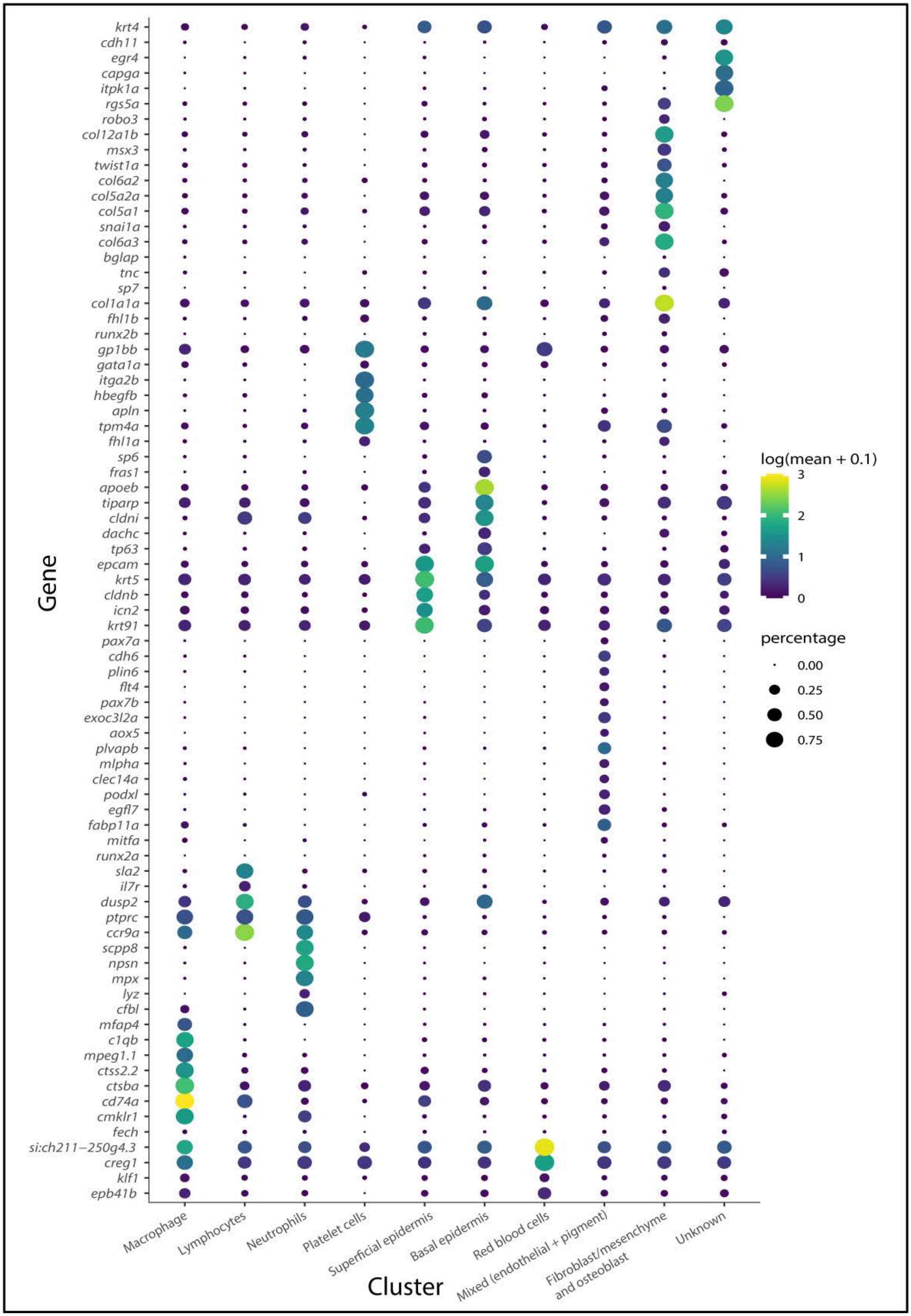


**Supplemental Figure 1**

**Figure S1. Marker gene expression used to define 7 day post amputation regenerating caudal fin single cell RNA-sequencing clusters.** Gene expression dot plots show transcript detection frequency (dot size) and average expression level (dot color) of established marker genes across the 10 Louvain clusters. These plots were used to infer cluster identities as follows: Red blood cells (2,744 cells), Fibroblast/mesenchyme and osteoblast (1,391 cells), Superficial epidermis (1,145 cells), Macrophage (722 cells), Lymphocytes (708), Basal epidermis (219 cells), Mixed (endothelial + pigment) (150 cells), Neutrophils (86 cells), Platelet cells (43 cells), and unknown (42 cells).

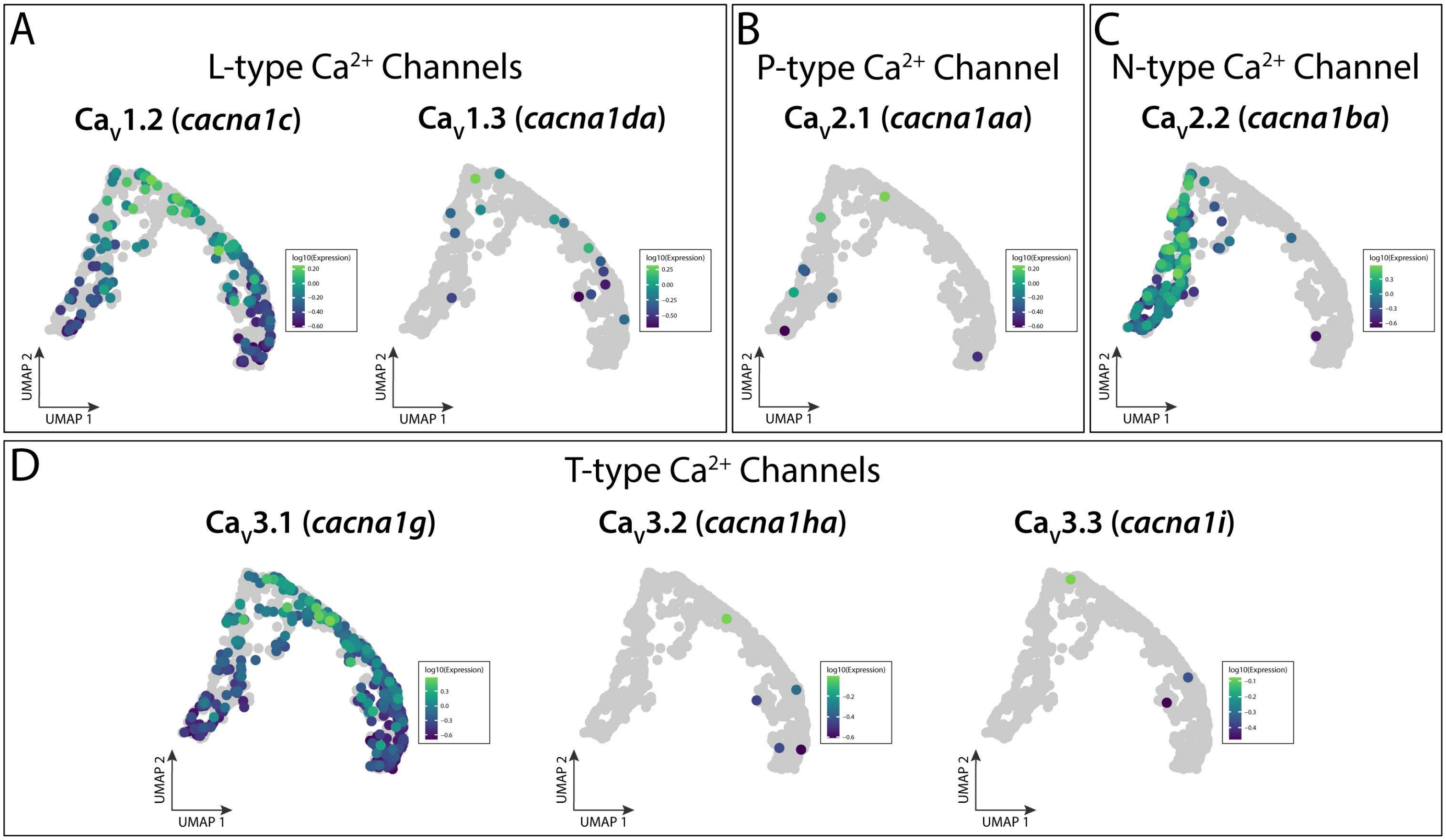


**Supplemental Figure 2**

**Figure S2. Identifying voltage-gated $Ca^{2+}$ channels expressed in the fibroblast/mesenchyme/osteoblast cluster. (A-D)** Candidate voltage-gated $Ca^{2+}$ channel types and UMAP visualization of their corresponding $\alpha_1$ subunit(s). Each panel represents L- **(A)**, P- **(B)**, N- **(C)**, or T-type **(D)** voltage-gated $Ca^{2+}$ channels. Only *cacna1c* (L-type), *cacna1ba* (N-type), and *cacna1g* (T-type) genes were predominantly expressed amongst fibroblast-lineage cells and/or osteoblast cells.

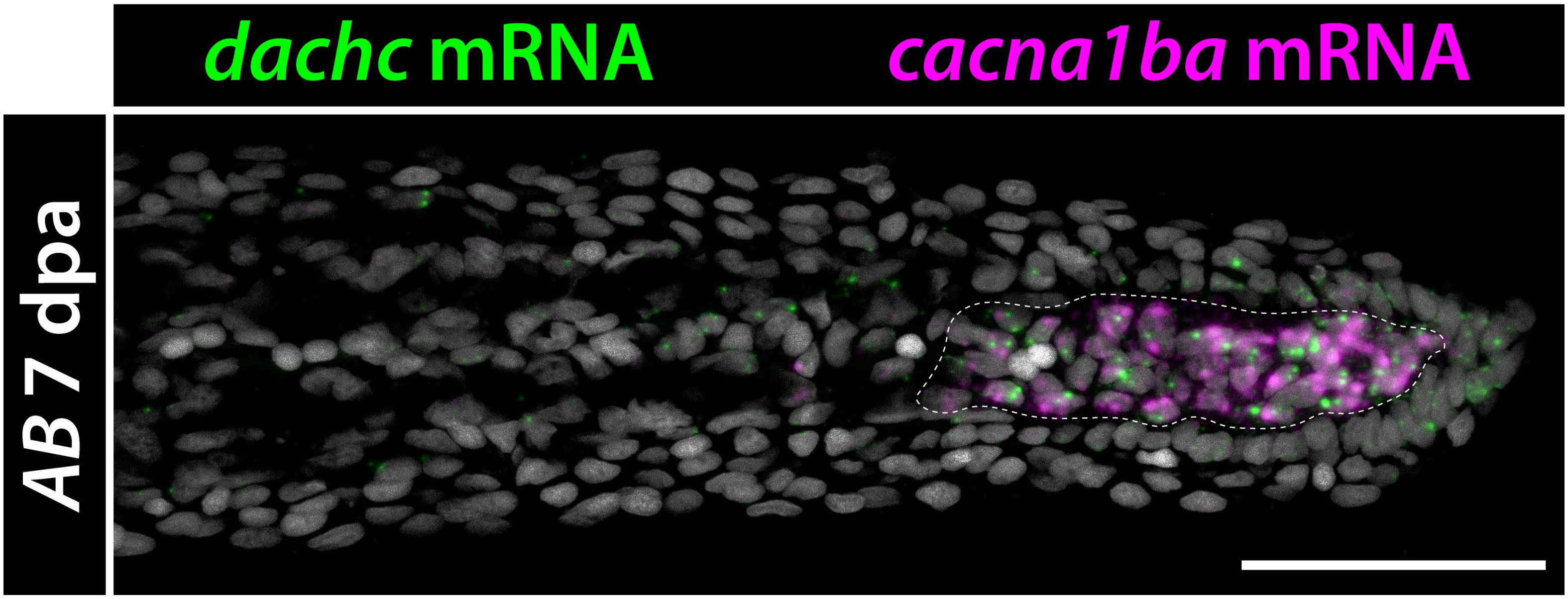


**Supplemental Figure 3**

**Figure S3.** ***cacna1ba*** **co-expresses with** ***dachshund c*** **(*****dachc*****) in distal blastema cells.** Confocal maximum intensity projection images showing double RNAscope in situ hybridization for *dachc* (green) and *cacna1ba* (magenta) mRNA on a 7 dpa wildtype regenerating caudal fin section. The dashed white line shows distalmost *cacna1ba* expression. Hoechst-stained nuclei are in gray. The scale bar is 50 µm.

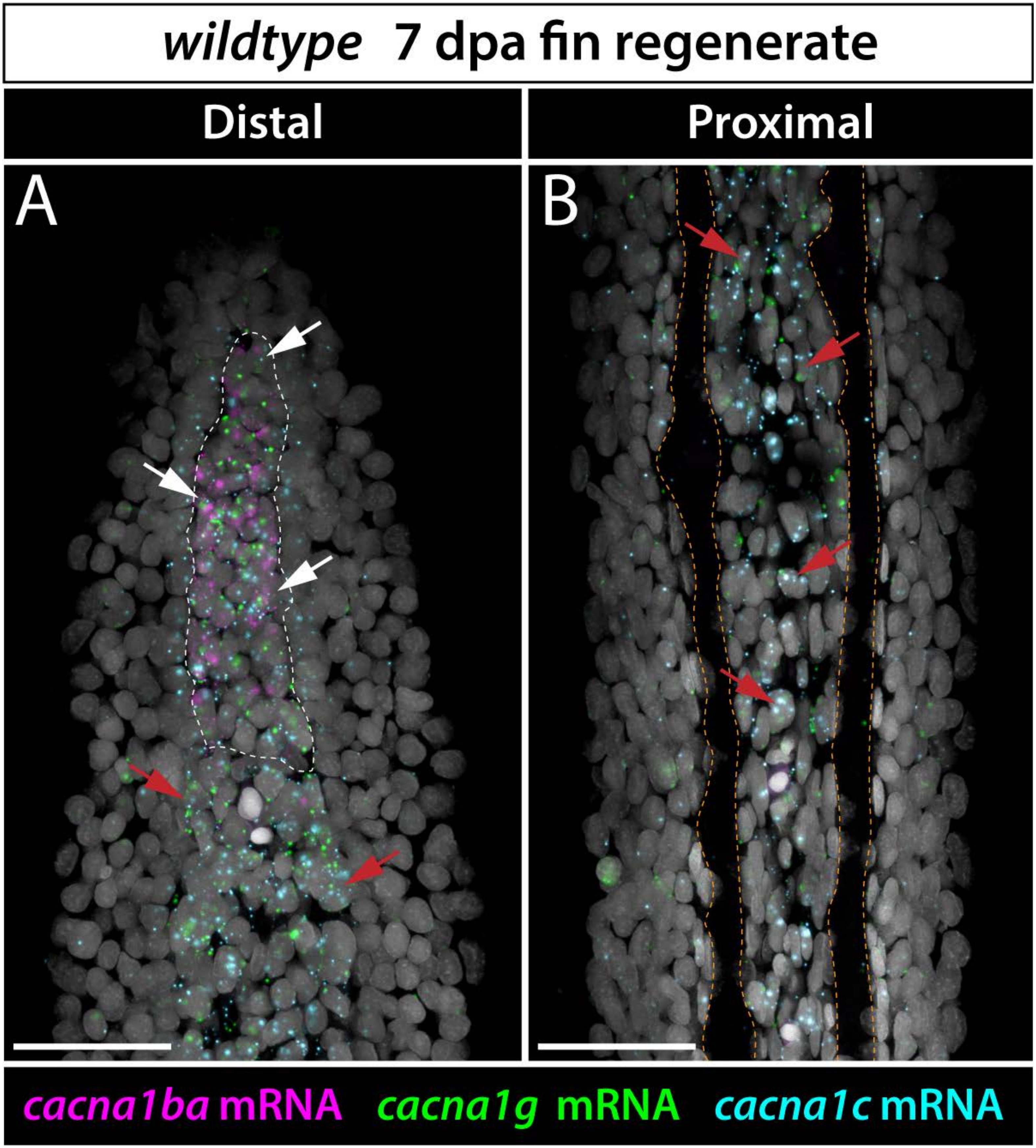


**Supplemental Figure 4**

**Figure S4. Co-expression of *cacna1c*, *cacna1ba*, and *cacna1g* in the distal blastema of regenerating fins.** Maximum intensity projection confocal images of *cacna1c*, *cacna1ba*, and *cacna1g* RNAscope staining at distal **(A)** and proximal **(B)** positions from Figure 3I. Dashed white region outlines distal fibroblast-lineage cells. White arrows point to cells co-expressing *cacna1c*, *cacna1ba*, and *cacna1g* mRNA and red arrows point to cells co-expressing *cacna1c* and *cacna1g* mRNA. Hoechst-stained nuclei are in gray. Dashed orange lines outline fin rays. Scale bars are 50 µm.

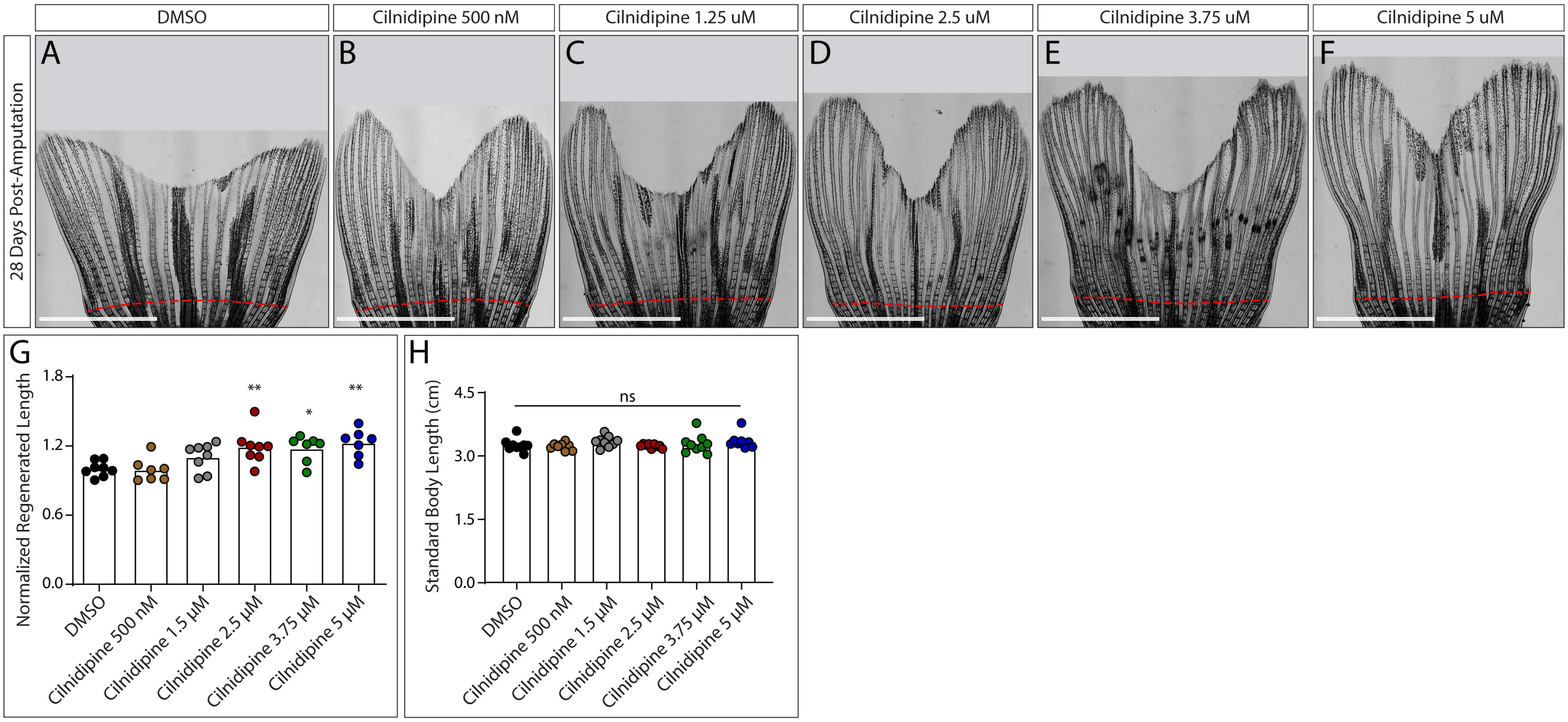


**Supplemental Figure 5**

**Figure S5. Zebrafish regenerate overgrown caudal fins across a range of cilnidipine doses. (A-F)** Stitched brightfield images of regenerated caudal fins after daily treatment with DMSO **(A)** or 500 nM **(B)**, 1.25 μM **(C)**, 2.5 μM **(D)**, 3.75 μM **(E)**, and 5 μM **(F)** cilnidipine from 5 – 28 dpa. The dashed red line indicates the amputation plane. The scale bars are 4 mm. **(G)** Graph showing regenerated caudal fin lengths (third ray) normalized to the mean of the DMSO-treated control group. Fish received daily 4-hour exposures of DMSO (n=8) or 500 nM (n=7), 1.25 μM (n=8), 2.5 μM (n=8), 3.75 μM (n=7), or 5 μM (n=7) cilnidipine. Each point represents an individual fish. **(H)** Graph showing standard body length measurements (tip of the mouth to the caudal peduncle) for adult wildtype treated with DMSO or cilnidipine at indicated dosages. *: $P < 0.05$ and **: $P < 0.01$ vs. the wildtype-DMSO treated group using one-way ANOVA and Tukey's post-hoc tests; ns: not significant.

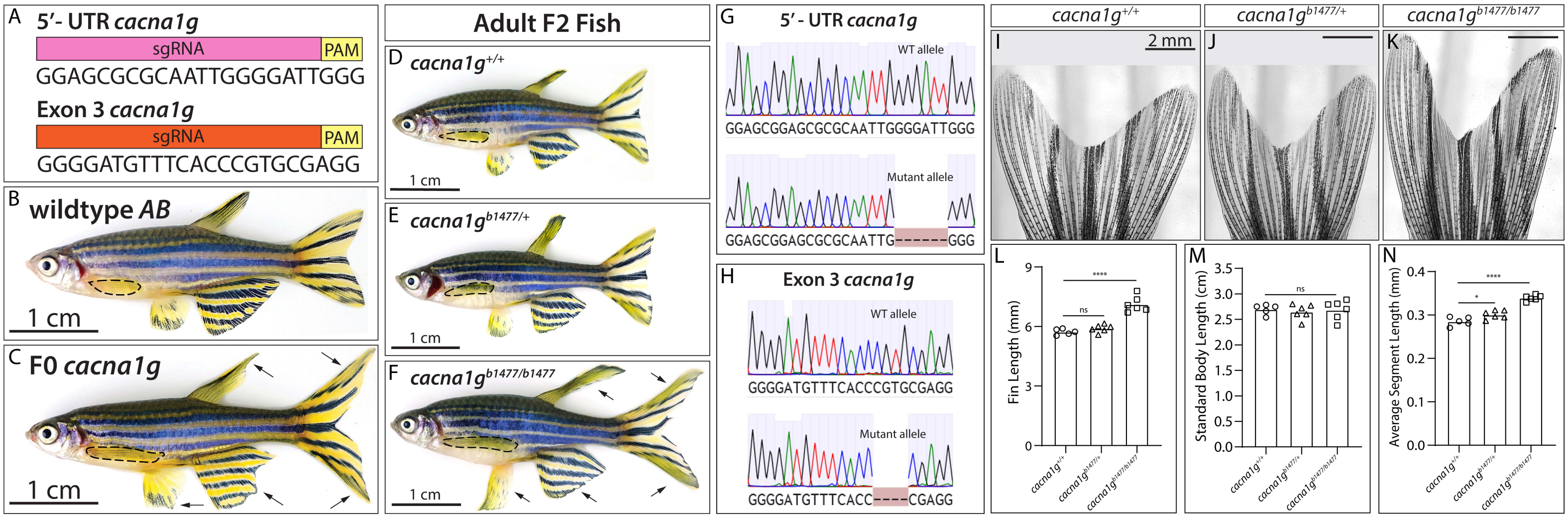


**Supplemental Figure 6**

**Figure S6.** ***cacna1g*** **homozygous loss-of-function modestly increases developed fin size. (A)** Sequence of the guide RNAs used to target genomic sequence in the 5′UTR and/or exon 3 of *cacna1g*. **(B, C)** Brightfield whole animal images of control wildtype *AB* **(B)** and F0 *cacna1g* crispant **(C)** fish. **(D-F)** Brightfield whole animal images of wildtype **(D)**, *cacna1g*$^{b1477/+}$ heterozygous **(E)**, and *cacna1g*$^{b1477/b1477}$ homozygous mutant **(F)** fish from the founder shown in panel C. Black arrows indicate overgrown tissue seen in all fins. Dashed black lines outline the pectoral fin. **(G, H)** Sanger sequencing chromatograms of PCR products from wildtype (WT) and mutant *cacna1g*$^{b1477}$ alleles showing a 6 bp deletion in the 5'UTR **(G)** and a 4 bp deletion in exon 3 **(H)**. **(I-K)** Brightfield stitched images of adult caudal fins from clutchmate *cacna1g*$^{+/+}$ **(I)**, *cacna1g*$^{b1477/+}$ **(J)**, and *cacna1g*$^{b1477/b1477}$ **(K)** animals. Scale bars are 2 mm. **(L)** Fin length measurements using the third ray (from a position perpendicular to the end of the longest procurrent ray to the fin tip). **(M)** Standard body length is unchanged with *cacna1g* loss-of-function. **(N)** Average ray segment length of the third caudal fin ray (n=5 segments/fish) of *cacna1g*$^{b1477/+}$ and *cacna1g*$^{b1477/b1477}$ animals. *: $P < 0.05$ and ****: $P < 0.0001$ vs. the wildtype group (*cacna1g*$^{+/+}$) using one-way ANOVA and Tukey's tests. ns: not significant.

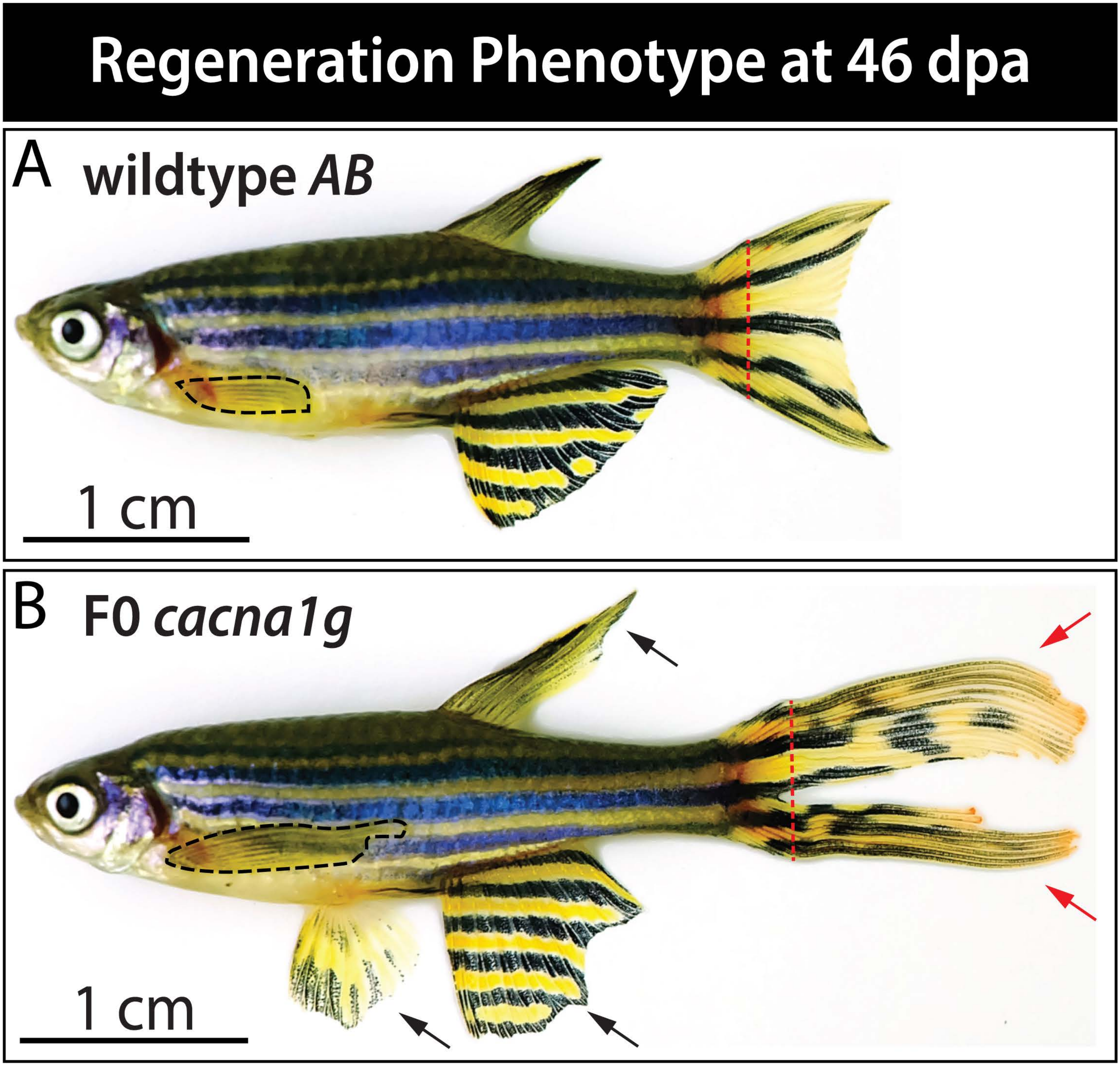


Supplemental Figure 7

**Figure S7. Developmental and regenerative fin overgrowth phenotype in *cacna1g* crispants.** **(A, B)** Brightfield images of a wildtype *AB* control and the *cacna1g* F0 crispant founder fish shown in Supplemental Figure 6 at 46 days post caudal fin amputation. The pelvic fin is not shown in **(A)**. Dashed black lines outline pectoral fins. Red dashed lines indicate the amputation plane. Black arrows point to regions of developmental fin overgrowth. Red arrows indicate dramatic regenerative caudal fin overgrowth. Scale bars are 1 cm.

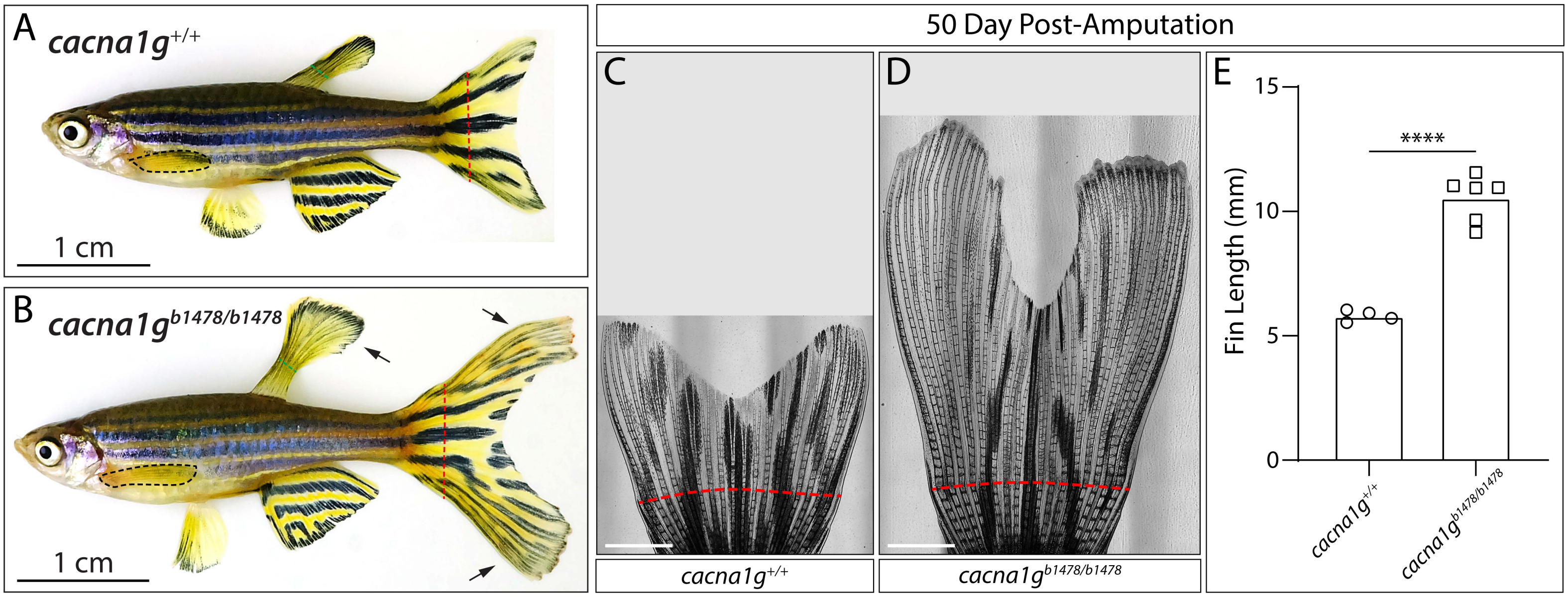


**Supplemental Figure 8**

**Figure S8. A second *cacna1g* loss-of-function allele recapitulates dramatic regenerative fin overgrowth.** Brightfield whole animal view of control wildtype **(A)** and homozygous *cacna1g*$^{b1478}$ mutant **(B)** zebrafish at 70 dpa of the caudal fin. The *cacna1g*$^{b1478}$ allele harbors a 4 bp deletion and a 36 bp insertion at the guide site that results in an early stop codon. Black dashed lines outline the pectoral fin. Green dashed lines mark where the dorsal fin was clipped for genotyping. Red dashed lines indicate caudal fin amputation site. Scale bars are 1 cm. **(C, D)** Brightfield stitched images of adult caudal fins at 50 dpa for *cacna1g*$^{+/+}$ **(C)** and *cacna1g*$^{b1478/b1478}$ **(D)**. Scale bars are 2 mm. **(E)** Caudal fin length measurements of *cacna1g*$^{+/+}$ and *cacna1g*$^{b1478/b1478}$ fish at 50 dpa. $^{****}$: $P<0.001$ by an unpaired Student's t-test.

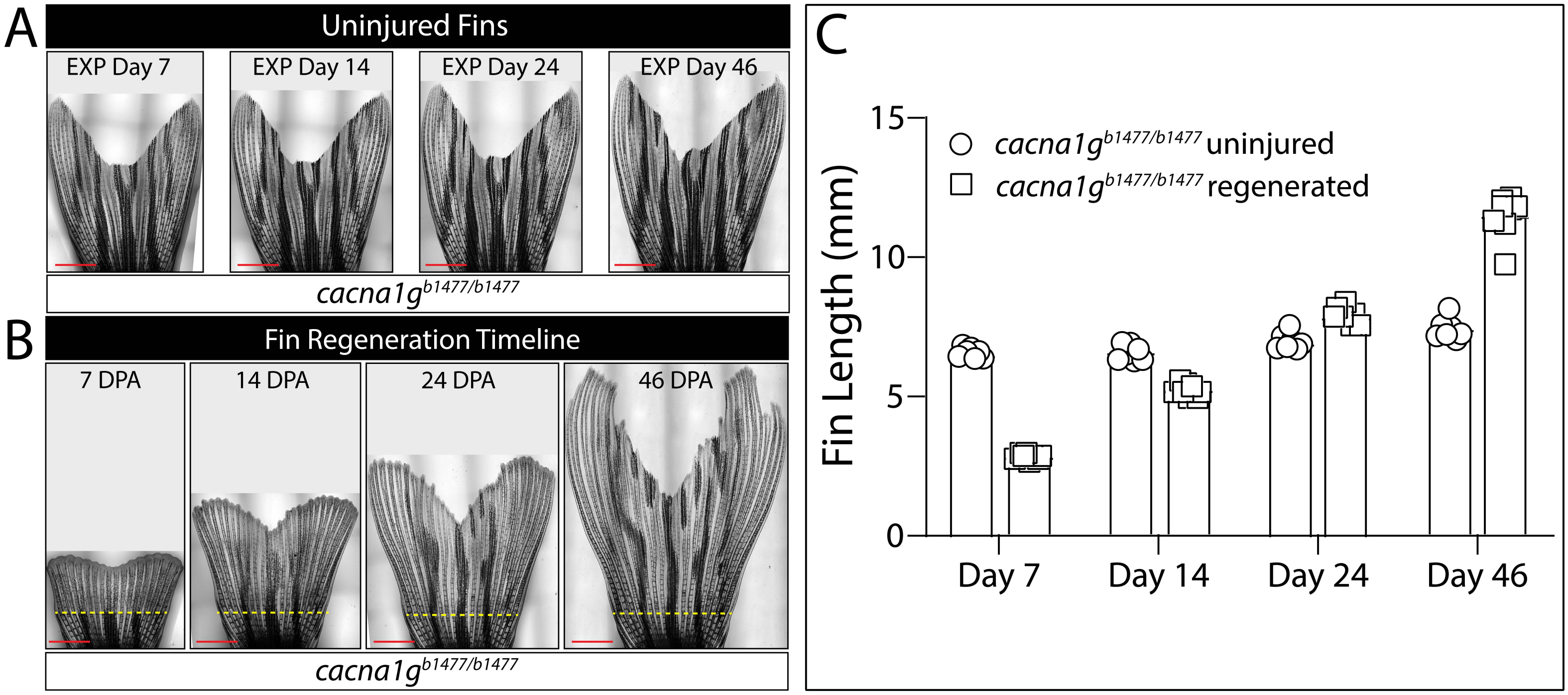


**Supplemental Figure 9**

**Figure S9. Cacna1g restores regenerating fins to their original scale. (A, B)** Brightfield stitched images of uninjured **(A)** and regenerating **(B)** *cacna1g*$^{b1477/b1477}$ caudal fins. Fin images were captured on days 7, 14, 24, and 46 of the experiment (abbreviated as EXP) or days post-amputation (dpa). Yellow dashed lines indicate the amputation plane. Red scale bars are 2 mm. **(C)** Graph showing caudal fin lengths of *cacna1g*$^{b1477/b1477}$ animals over the indicated time course from groups left uninjured (circles) or amputated (squares).

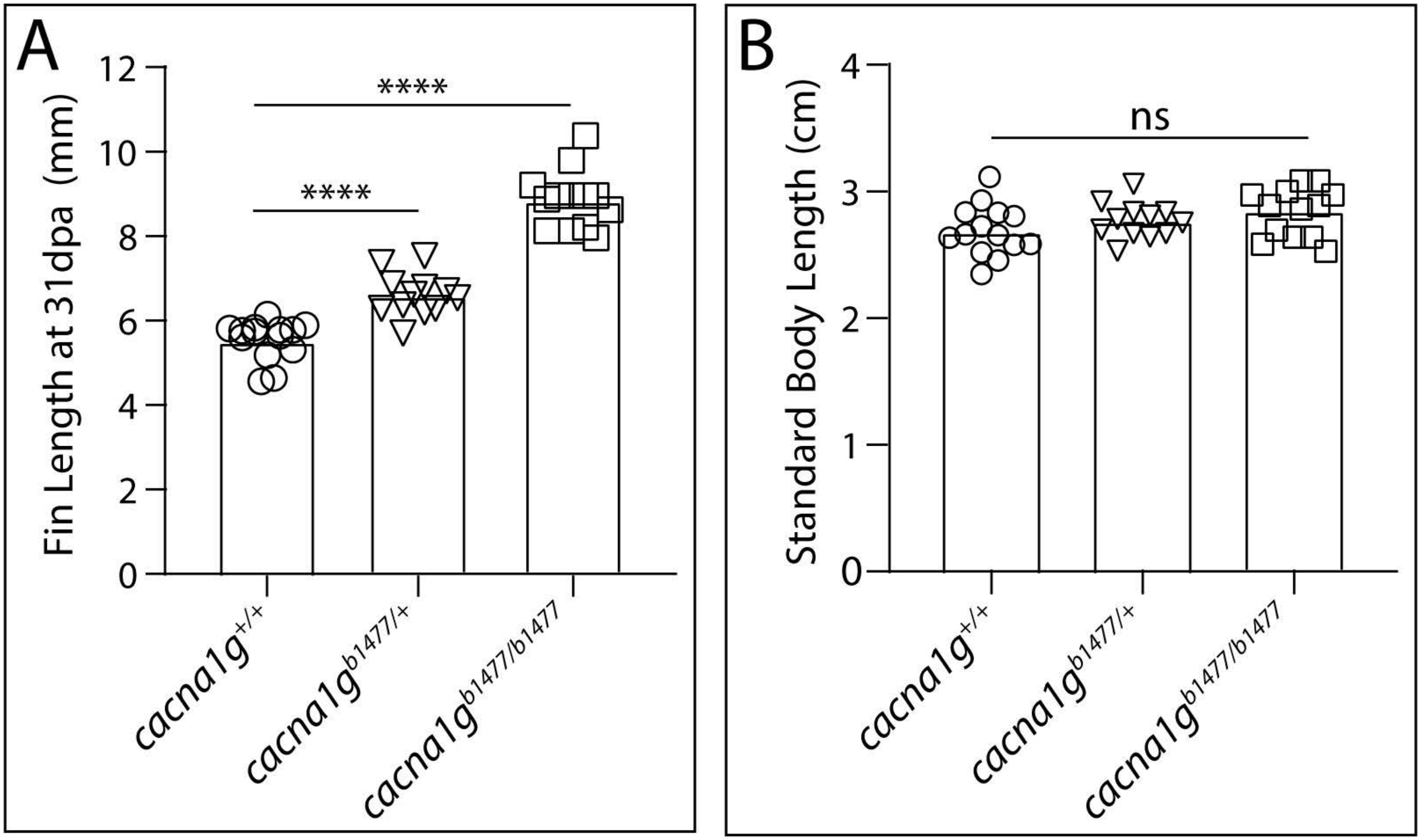


**Supplemental Figure 10**

**Figure S10.** ***cacna1g* heterozygotes display mild fin regenerative overgrowth. (A, B)** Graphs showing regenerated caudal fin lengths **(A)** and standard body lengths **(B)** at 31 dpa for *cacna1g*$^{+/+}$ (circles), *cacna1g*$^{b1477/+}$ (triangles), and *cacna1g*$^{b1477/b1477}$ (squares) fish. Each data point represents an individual animal (n=14 fish per genotype). ****: $P > 0.0001$ using one-way ANOVA and Tukey's tests; ns: not significant.

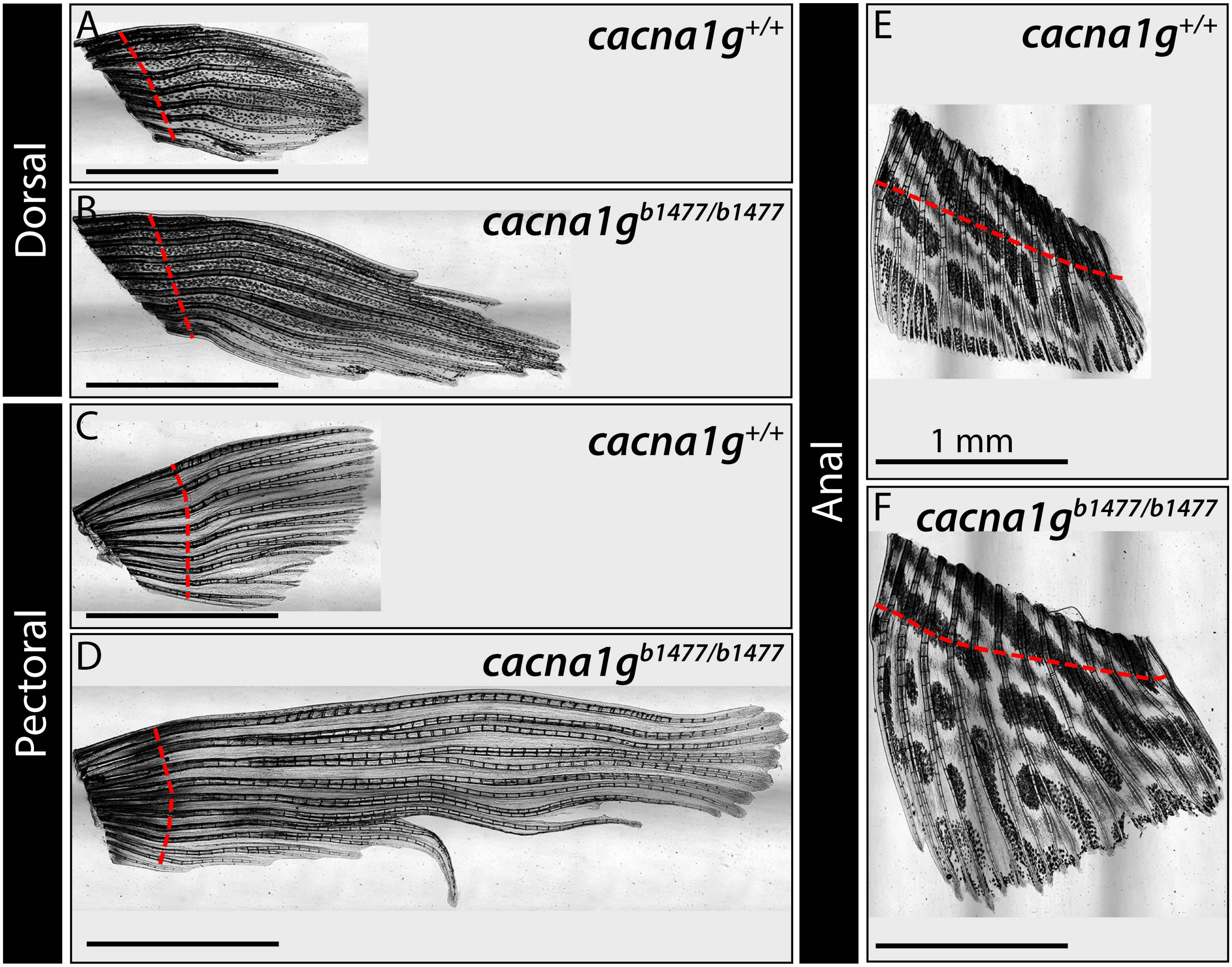


**Supplemental Figure 11**

**Figure S11. All median and paired fins of *cacna1g*$^{b1477/b1477}$ fish regenerate to extraordinary length. (A-F)** Brightfield stitched images contrasting regenerated fins dissected from *cacna1g*$^{+/+}$ and *cacna1g*$^{b1477/b1477}$ fish at 46 dpa. Representative dorsal fins **(A, B)**, pectoral fins **(C, D)**, and anal fins **(E, F)** are shown. Pelvic fins are not shown but behave likewise. Dashed red lines indicate amputation positions. Scale bars are 1 mm.

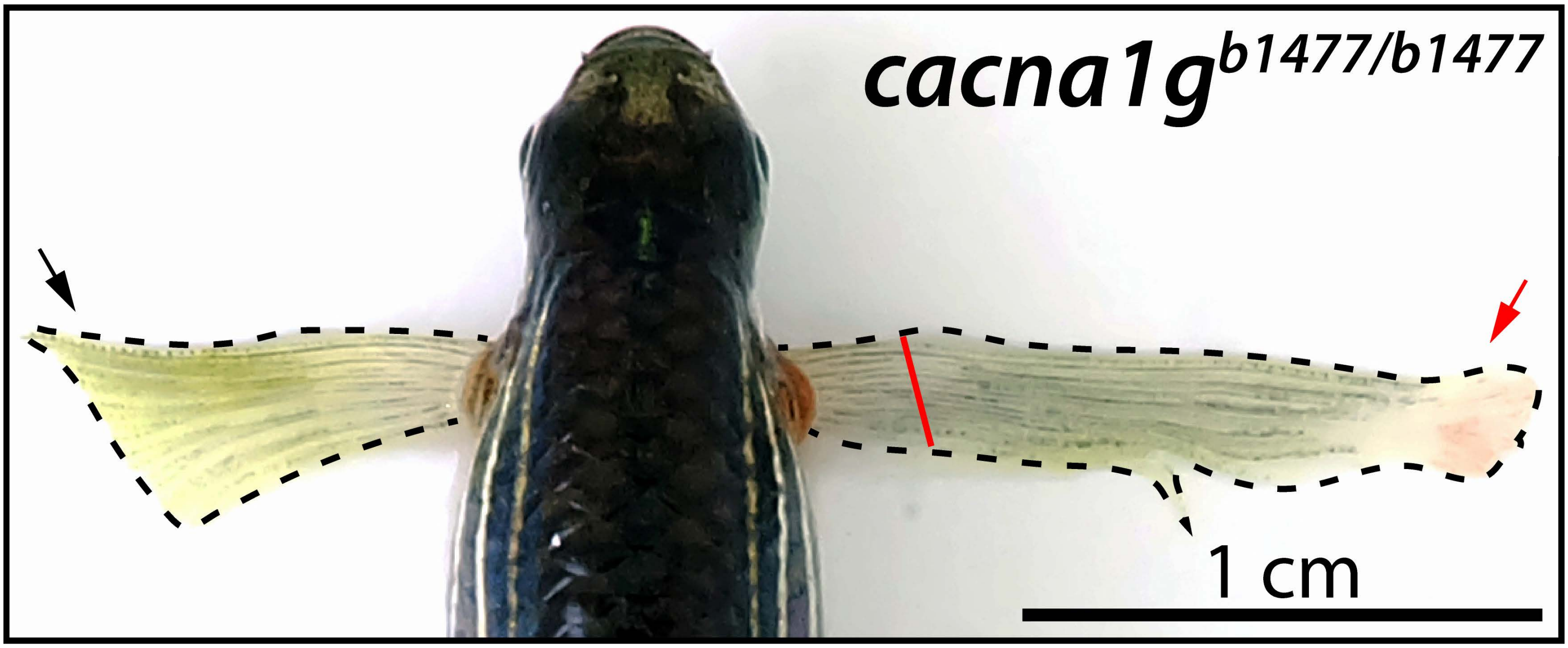


**Supplemental Figure 12**

**Figure S12. Unilateral pectoral fin overgrowth reinforces that Cacna1g relatively impacts regenerative over developmental outgrowth.** Brightfield dorsal view image of a *cacna1g*$^{b1477/b1477}$ adult at 46 days post amputation of the right pectoral fin. Black dashed lines outline the pectoral fins. The red line marks the amputation site. The red arrow indicates dramatically overgrown tissue of the regenerating pectoral fin. The black arrow shows the milder, developmental outgrowth in the uninjured paired fin. The scale bar is 1 cm.

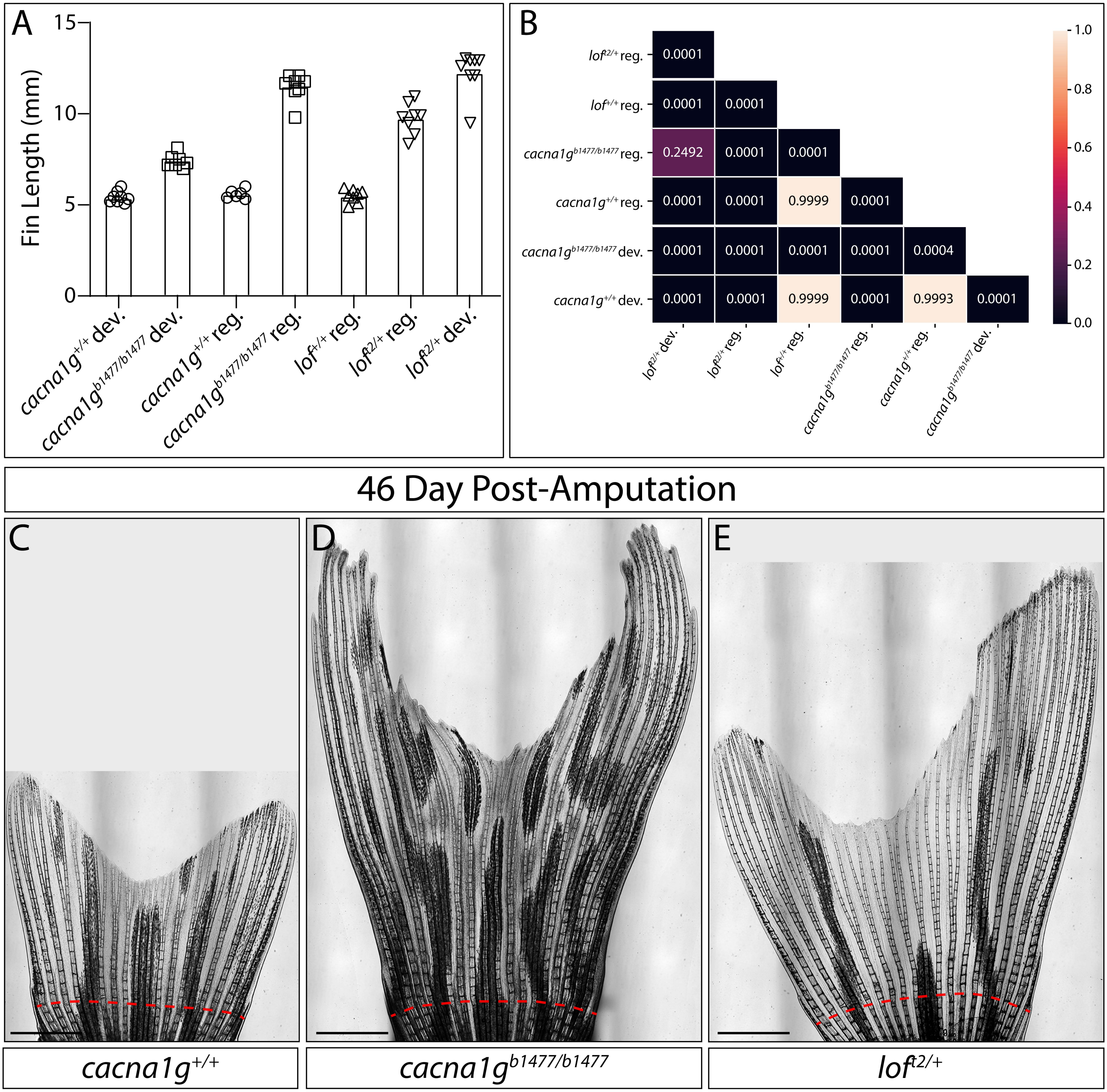


**Supplemental Figure 13**

**Figure S13. The *cacna1c* loss-of-function regenerated long-finned phenotype exceeds that of *longfin*$^{t2}$ heterozygotes. (A)** Graph comparing developed and regenerated caudal fin lengths of *cacna1g*$^{b1477/b1477}$ and *longfin*$^{t2/+}$ fish along with corresponding clutchmate controls. **(B)** Heatmap depicting P-values for all cross-comparison of the groups in **(A)**, following a one-way ANOVA and Tukey's tests. **(C-E)** Brightfield stitched images of *cacna1g*$^{+/+}$ **(C)**, *cacna1g*$^{b1477/b1477}$ **(D)**, and *longfin*$^{t2/+}$ **(E)** caudal fins at 46 dpa. Red dashed lines indicate amputation planes. Scale bars are 1 mm.

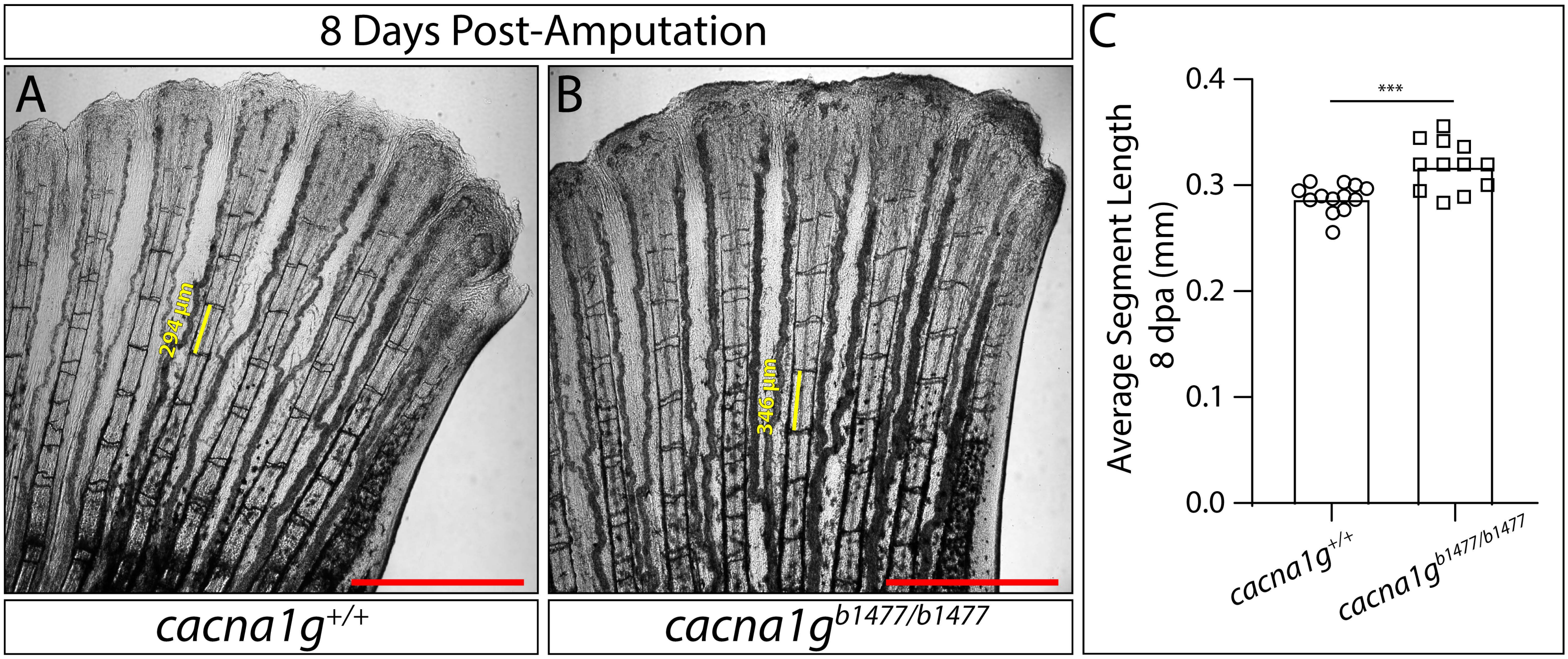


**Supplemental Figure 14**

**Figure S14. Overgrown *cacna1g*$^{b1477/b1477}$ regenerated caudal fins retain joints with near-normal spacing. (A, B)** Brightfield stitched images of the caudal fin ventral lobe of *cacna1g*$^{+/+}$ **(A)** and *cacna1g*$^{b1477/b1477}$ **(B)** fish at 8 days post amputation (dpa). Yellow lines indicate example individual segment lengths. Scale bars are 1 mm. **(C)** Graph showing the average ray segment length (n=12 or 13 fish, each data point is the mean of 5 segments) of 8 dpa regenerated caudal fins from *cacna1g*$^{+/+}$ and *cacna1g*$^{b1477/b1477}$ fish. ***: $P<0.001$ vs. the control group (*cacna1g*$^{+/+}$) using an unpaired Student's t-test.

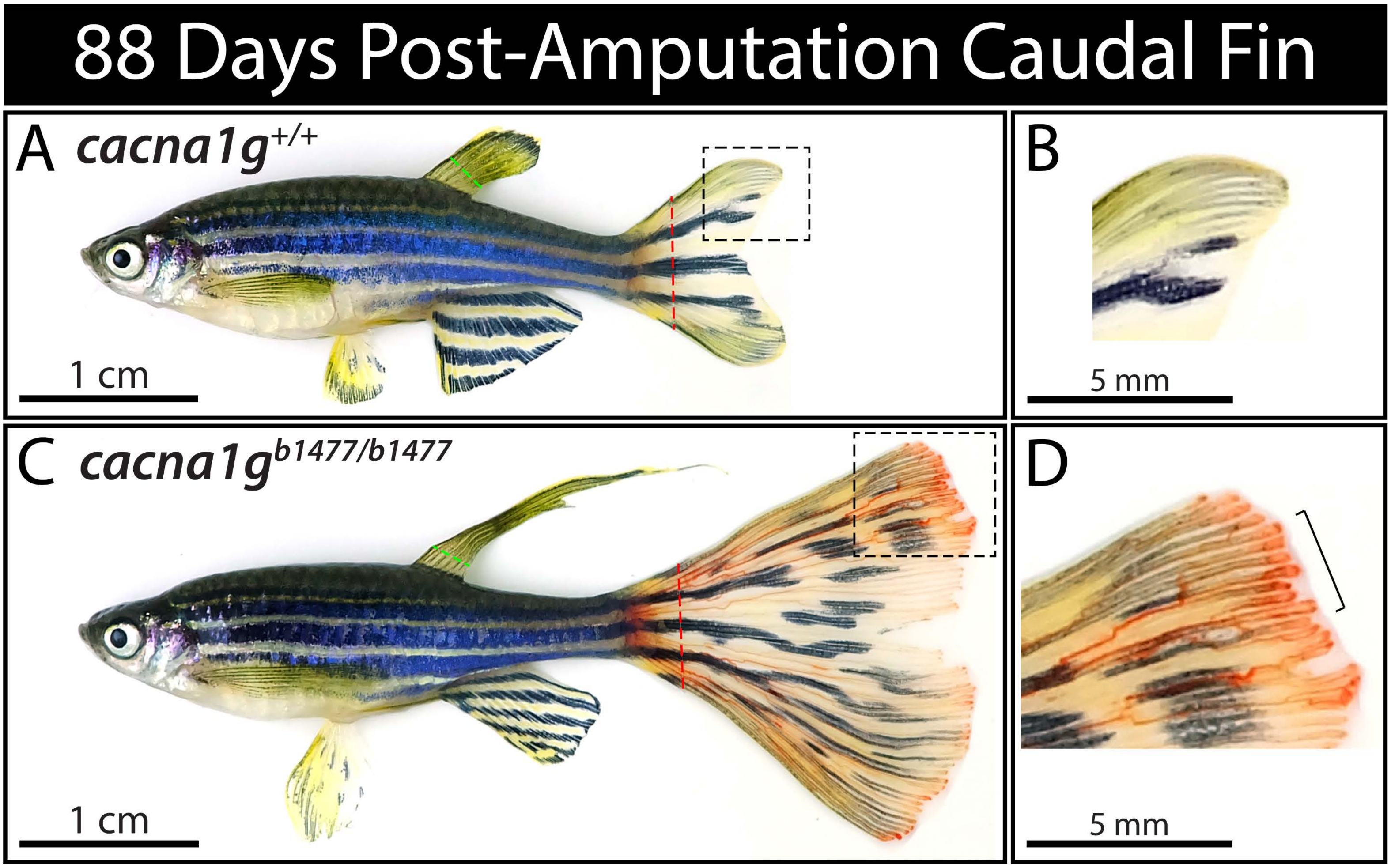


**Supplemental Figure 15**

**Figure S15. Blood pooling in *cacna1g*$^{b1477/b1477}$ regenerating adult caudal fins. (A-D)** Brightfield images of *cacna1g*$^{+/+}$ **(A, B)** and *cacna1g*$^{b1477/b1477}$ **(C, D)** fish at 88 days post caudal fin amputation. Dashed green lines mark dorsal fin clipping positions (for genotyping). Red dashed lines indicate caudal fin amputation planes. Black dashed boxes outline the zoomed-in distal caudal fin regions shown in **(B)** and **(D)**. The black bracket highlights a region of blood pooling. Scale bars are 1 cm or 5 mm.

**Movie S1. Blastemal cells of regenerating caudal fins exhibit dynamic $Ca^{2+}$ transients.** Confocal time-lapse imaging of the caudal fin distal blastema from a live *tph1b:GCaMP6s* fish at 3 days post amputation (dpa). GCaMP6s fluorescence is green. Elapsed time is in minutes and seconds. Arrows indicate representative cells with dynamic GCaMP6s fluorescence. The scale bar is 20 μm.

**Movie S2. Isolated regenerating fin fibroblasts spontaneously flux $Ca^{2+}$ and respond to depolarization.** Time-lapse confocal images of primary fin cells isolated from *tph1b:GCaMP6s* 5 dpa regenerating caudal fins. Green is GCaMP6s fluorescence. Membrane depolarization by KCl addition to a final concentration of 80 mM occurs at 2 minutes and 6 seconds. Ionomycin treatment at 5 minutes and 5 seconds serves as a positive control for GCaMP6s responsiveness and to calibrate each cell's maximum GCaMP6s fluorescence. Elapsed time is in minutes and seconds. The scale bar is 50 μM.

**Movie S3. Cacna1g enables extensive and dynamic $Ca^{2+}$ fluxes within the distal blastema during regenerative fin outgrowth.** Confocal GCaMP6s (green; *tph1b:GCaMP6s*) time-lapse images of caudal fin distal blastemas of $cacna1g^{+/+}$ and $cacna1g^{b1477/b1477}$ fish at 5 days post amputation. Elapsed time is in minutes and seconds. White arrows indicate $Ca^{2+}$-fluxing cells. The scale bar is 30 μm.

`
